# Native size exclusion chromatography-based mass spectrometry (SEC-MS) identifies novel components of the Heat Shock Protein 90-dependent proteome

**DOI:** 10.1101/2022.05.23.492985

**Authors:** Rahul S. Samant, Silvia Batista, Mark Larance, Bugra Ozer, Christopher I. Milton, Isabell Bludau, Laura Biggins, Simon Andrews, Alexia Hervieu, Harvey E. Johnston, Bissan Al-Lazikhani, Angus I. Lamond, Paul A. Clarke, Paul Workman

## Abstract

The molecular chaperone heat shock protein 90 (HSP90) works in concert with co-chaperones to stabilize its client proteins, which include multiple drivers of oncogenesis and malignant progression. Pharmacologic inhibitors of HSP90 have been observed to exert a wide range of effects on the proteome, including depletion of client proteins, induction of heat shock proteins, dissociation of co-chaperones from HSP90, disruption of client protein signaling networks, and recruitment of the protein ubiquitylation and degradation machinery—suggesting widespread remodeling of cellular protein complexes. However, proteomics studies to date have focused on inhibitor-induced changes in total protein levels, often overlooking protein complex alterations. Here, we use size-exclusion chromatography in combination with mass spectrometry (SEC-MS) to characterize the changes in native protein complexes following treatment with the HSP90 inhibitor tanespimycin (17-AAG) in the HT29 colon adenocarcinoma cell line. After confirming the signature cellular response to HSP90 inhibition (e.g., induction of heat shock proteins, decreased total levels of client proteins), we were surprised to find only modest perturbations to the global distribution of protein elution profiles in inhibitor-treated cells. Similarly, co-chaperones that co-eluted with HSP90 displayed no clear difference between control and treated conditions. However, two distinct analysis strategies identified multiple inhibitor-induced changes, including several known components of the HSP90 proteome, as well as numerous proteins and protein complexes with no previous links to HSP90. We present this dataset as a resource for the HSP90, proteostasis, and cancer communities (https://www.bioinformatics.babraham.ac.uk/shiny/HSP90/SEC-MS/), laying the groundwork for future mechanistic and therapeutic studies related to HSP90 pharmacology. Data are available via ProteomeXchange with identifier PXD033459.

## Introduction

The molecular chaperone Heat Shock Protein 90 (HSP90) is required for the stabilization and activation of around 300 client proteins (see http://www.picard.ch/downloads/Hsp90interactors.pdf for the latest client list), many of which are oncogenic kinases that are mutated and/or hyper-activated in malignancies (Jaeger and Whitesell, 2019). Furthermore, HSP90 may act as an ‘enabler’ of oncogenesis and malignant progression, potentially supporting tumor heterogeneity and contributing to drug resistance (Lacey and Lacey, 2021). Pharmacologic inhibitors of HSP90 have therefore been pursued as anti-cancer agents, either alone or in combination. However, none of the 18 HSP90 inhibitors clinically tested to date have shown sufficient efficacy and tolerability to progress to FDA approval (Yuno et al., 2018; Workman, 2020). Part of the discrepancy between promising *in vitro* data and underwhelming clinical benefit could be due to insufficient data on the pharmacodynamic response to HSP90 inhibition (Butler et al., 2015). Nevertheless, there is continuing interest in HSP90 as a pharmacologic cancer target, including in anti-cancer immunotherapy (Zavareh et al., 2021), as well as a recent application for approval of one HSP90 inhibitor for chemotherapy-relapsed gastro-intestinal stromal tumors (https://www.taiho.co.jp/en/release/2021/20210914.html)(Honma et al., 2021)). HSP90 inhibition has also shown potential beyond the cancer field, e.g., as a broad-spectrum anti-viral (Wang et al., 2017) (including activity against SARS-CoV2 (Goswami et al., 2021)), and as a gero-protector for healthier ageing (Fuhrmann-Stroissnigg et al., 2018; Janssens et al., 2019)). Therefore, improving our understanding of the molecular responses to HSP90 inhibition at a global, proteome-wide scale could provide more rational strategies for patient selection and stratification across a variety of pathologies.

Global approaches could also help clarify the mechanisms underlying the tumor selectivity of HSP90 inhibitors—a phenomenon that has long been a matter of debate (Chiosis and Neckers, 2006; Workman, 2004). HSP90 functions as a homodimer and activates its client proteins in an ATP-dependent manner in concert with dozens of co-chaperones (Pearl, 2016; Schopf et al., 2017). Initially reported for the oncogenic protein kinase clients ERBB2 and CRAF in cancer cell lines treated with the natural product HSP90 inhibitor geldanamycin (Mimnaugh et al., 1996; Schulte et al., 1995, 1996), it is now well-established that HSP90–client protein complex disruption, client ubiquitylation, and client degradation by the proteasome are all general components of the cellular response to pharmacologic HSP90 inhibition. Given the number of oncoproteins that make up the HSP90 client list, a major rationale for deploying HSP90 inhibitors in cancer is the destabilisation and disruption of signalling networks critical for oncogenesis and malignant progression. However, several alternative or additional mechanisms have been proposed, including the tumor-selective accumulation of several different HSP90 inhibitors, as well as their much higher affinity for the hyper-activated HSP90 complexes specifically observed in tumor cells (Chiosis and Neckers, 2006; Kamal et al., 2003; Wang et al., 2019). This latter stress-associated assembly of high molecular-weight complexes, containing multiple chaperones and co-chaperones— more recently referred to as the ‘epichaperome’—may potentially be more predictive of patient response than expression levels of the chaperones or their individual oncogenic protein clients *per se* (Wang et al., 2019). Furthermore, systemic dysregulation of chaperones and other quality control machineries that together make up the proteostasis network is a hallmark of several other ageing-related pathologies (Hipp et al., 2019). Given the clinical interest in exploiting disease-altered states of epichaperome and proteostasis networks (Brehme et al., 2019), unbiased systems-wide analysis of how such higher-order protein assemblies are perturbed by HSP90 inhibitors and other proteostasis modulating agents could be valuable for maximising therapeutic benefit.

To date, proteomic studies aiming to characterize the HSP90-dependent proteome can be separated broadly into two categories (Weidenauer et al., 2017). The first set of these comprise mass spectrometry-based comparative proteomics to identify proteins whose abundances change following HSP90 inhibitor treatment (Maloney et al., 2007; Schumacher et al., 2007; Sharma et al., 2012; Voruganti et al., 2013; Wu et al., 2012). The experimental design of such HSP90 inhibition-altered proteome studies generally involves use of inhibitor concentrations and/or time-points that result in degradation of well-characterized client proteins. While this approach undoubtedly has been fruitful, it misses client proteins whose levels do not change drastically, have slower degradation kinetics, or form non-functional oligomers/aggregates. It also ignores functionally consequential alterations in protein complexes whose total abundances would not be expected to change, including components of epichaperome assemblies (e.g., co-chaperones, ubiquitylation enzymes), as well as a diverse range of protein complexes reliant on HSP90 for their correct assembly and maintenance (Makhnevych and Houry, 2012). To highlight this last point, a recent multi-parametric study found that HSP90 inhibition elicits much more widespread alterations to the proteome on the basis of changes in protein solubility, rather than changes in total abundance, across the same biological samples (Sui et al., 2020). Notably, only the solubility-based comparisons could identify protein complex subunits known to require HSP90 for their assembly into mature complexes but not for their stability.

Some of the limitations inherent in abundance-based differential proteomics can be addressed through the second category of proteomics approaches, which employ direct ‘interactomics’ combining affinity-based assays with mass spectrometry (Rodina et al., 2016) or high-content fluorimetry (Taipale et al., 2012) as a readout (Weidenauer et al., 2017). However, these bait : prey-based techniques are also limited in scope. They are mostly unsuitable for detecting weak or highly labile interactions, and changes observed in a protein : protein interaction following inhibitor treatment could be confounded by inhibitor-induced changes in the protein’s total abundance—thus making it difficult to interpret the data without extensive follow-up validation. Furthermore, indirect or downstream effects on protein complexes that do not interact with the HSP90 machinery are ignored.

Ideally, studies would incorporate the strengths of both approaches, allowing global identification of proteins that change in absolute abundance and/or in their distribution across different protein complexes—all in a single experiment. One potential solution is native SEC-MS, first employing size-exclusion chromatography (SEC) to separate protein complexes from a cell homogenate (lysed under non-denaturing conditions) into different fractions according to their molecular weight, followed by bottom-up mass spectrometry (MS) of each individual fraction (Heusel et al., 2019; Kirkwood et al., 2013; Salas et al., 2020). Importantly, SEC-MS allows analysis of endogenous protein complexes in cells without having to rely on affinity-tagged bait and/or overexpression systems, which have the potential to introduce artefacts. In this way, SEC-MS provides native molecular weight-based elution profiles together with total abundance at a proteome-wide level.

Here, we performed SEC-MS to characterize global changes to native protein complex distributions upon HSP90 inhibition with the geldanamycin-derivative tanespimycin (17-AAG) in the HT29 human colon adenocarcinoma cell line. We chose a tanespimycin concentration (62.5 nm) demonstrated to trigger the molecular signature of HSP90 inhibition (e.g., HSP70 induction) in this cell line, but at an early enough treatment time (8 h) that the majority of client degradation had yet to take place, so that we would be able to see remodeling of client-containing protein complexes (Samant et al., 2014). We identified 6,427 unique proteins overall, including 4,645 in at least three of the four biological replicates. Known members of well-characterized protein complexes displayed similar SEC-MS elution profiles. We were surprised to find minimal changes to the profiles of most identified proteins following HSP90 inhibition—including co-chaperones that dissociated from HSP90 clients under the same treatment conditions in previous immunoprecipitation studies. The lack of changes to co-chaperones detected by SEC-MS—which was confirmed by independent SEC-Immunoblots—was not due to a lack of target engagement, as the molecular signature of HSP90 inhibition was observed throughout our experiments. Nevertheless, we used two distinct analysis strategies to identify proteins and protein complexes whose SEC-MS profiles changed robustly in our dataset. These included several proteins previously characterized as being HSP90-dependent, as well as numerous novel hits—two of which we validated for biological importance. We present this dataset as a resource to the HSP90, proteostasis, and cancer communities (available to explore as a web-based Shiny app at https://www.bioinformatics.babraham.ac.uk/shiny/HSP90/SEC-MS/), providing novel candidates for further mechanistic and therapeutic studies.

## Results

### SEC-MS provides a platform to investigate changes in protein complex distributions following HSP90 inhibition

We treated HT29 colon adenocarcinoma cells with the HSP90 inhibitor tanespimycin (HSP90i), or mock-treated with DMSO vehicle (Control), for 8 hours before lysing in phosphate-buffer saline (PBS) without detergents or any other chaotropes that disrupt protein–protein interactions (Fig 1A). At this tanespimycin exposure concentration and time, we had previously shown that client protein interactions with HSP90 co-chaperones (e.g., CDC37, STIP1/HOP, AHA1) are disrupted but major client degradation has yet to occur (Samant et al., 2014). Lysates from each of the four Control or HSP90i replicates (eight samples total) were fractionated by SEC to separate the protein complexes by molecular weight into 24 sequential fractions of equal volume (Fig 1B). Each fraction was then subjected to standard protocols for bottom-up proteomics involving enzymatic digestion and LC-MS/MS, as described previously (Kirkwood et al., 2013). To improve proteome coverage, we divided each fraction in two and digested one with a 1:1 mixture of the proteases LysC and trypsin, and the other with trypsin alone. Thus, we generated a total of 384 samples for LC-MS/MS analysis. The resulting raw data were analyzed using MaxQuant software (Tyanova et al., 2016). Overall, MaxQuant identified 111,365 peptides across 7,401 proteinGroups (Table S1). After removing false identifications, and using a threshold of at least two peptides detected per protein at a false discovery rate (FDR) < 0.01, this equated to 6,804 protein groups, representing 6,427 unique proteins (following consolidation of duplicates into single entries)(Tables S2–S3).

**Figure 1.**
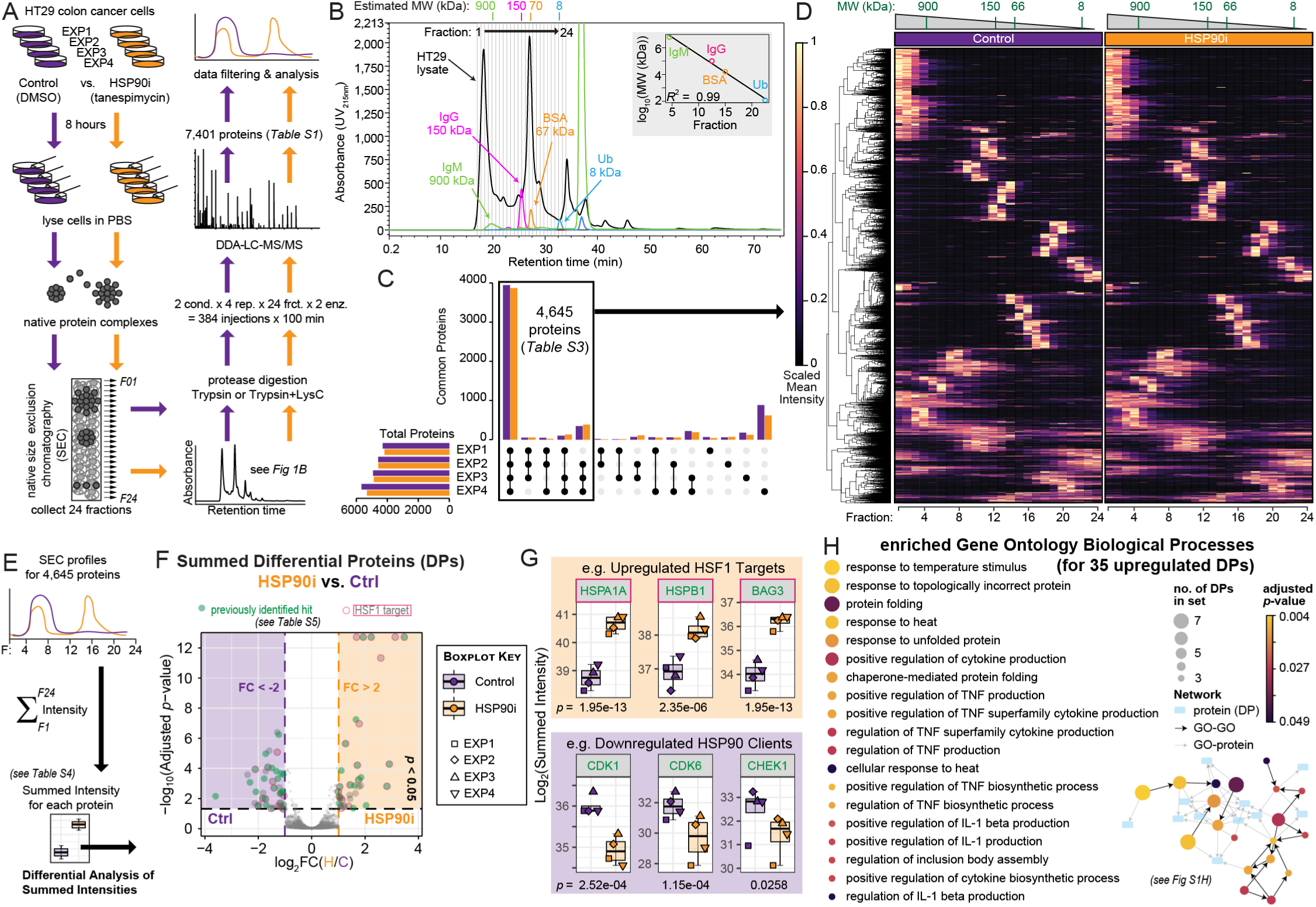
SEC-MS approach to investigate changes in native protein complex distributions following HSP90 inhibition. (**A**) Workflow for SEC-based protein complex isolation and LC-MS/MS-based identification in HT29 colon cancer cells treated with 5 x GI_50_ (62.5 nM) of the HSP90 inhibitor tanespimycin or mock-treated with DMSO. (**B**) UV chromatogram from one of the four control (DMSO-treated) biological replicates displaying the profile of the HT29 total cell lysate as it eluted from the Superose® 6 SEC column across 24 fractions. The retention time (in minutes) and UV absorbance (at 215 nm) are represented on the *x* - and *y* - axes, respectively. Protein standards of known molecular weights (IgM, Immunoglobulin M; IgG, Immunoglobulin G; BSA, Bovine Serum Albumin; Ub, Ubiquitin) were injected onto the same column, and their elution peaks were used to estimate the molecular weight (MW) range for each fraction using the *R* package *CCprofiler*. (**C**) Upset plot showing number of proteins identified in each of the four biological replicates (Control, purple, and HSP90i, orange, represented separately). Same data depicted as Venn Diagrams in Fig S1A–B. (**D**) Heatmap of scaled mean intensities for each of the 4,645 proteins filtered in (C). Mean intensities across the four replicates were calculated for each protein by fraction (1–24) and condition (Control or HSP90i). Scaling was performed across all 48 fractions, such that the highest fraction intensity value for a protein was set at 1 regardless of whether it was observed in the Control or HSP90i condition. The dendogram cut-offs based on Euclidean distance matrix with the Ward-D2 linkage method are illustrated to the left of the heat map. (**E**) Approach to calculate differentially-abundant proteins based on summed intensities across all 24 SEC fractions, using the *R* package DEP. See also Fig S1I and Table S5. (**F**) Volcano Plot calculated using the *R* package DEP, based on summed intensities for each of the 4,645 filtered proteins. Log_2_ -transformed Fold Changes (log_2_ FC) and negative log_10_ -transformed adjusted *p*-values (two-tailed Student’s *t* -test with Benjamini-Hochberg correction) are plotted on the *x* - and *y* - axes, respectively. Proteins with *p* < 0.05 and absolute log_2_ FC > 1 (i.e., FC > 2) are magnified. Proteins identified as ‘hits’ in previous high-throughput HSP90 proteomics studies are in green, and HSF1 targets are outlined in magenta (see also Table S6). (**G**) Summed intensities of known protein products of HSF1-activated genes (orange) and HSP90 clients (purple) are increased and decreased, respectively, following tanespimycin treatment. Box-and-whisker (Tukey) plots represent median, interquartile range, and absolute range for the four biological replicates. Adjusted *p*-values (Benjamini-Hochberg correction) calculated during differential expression analysis in (E) are indicated below each plot. (**H**) Gene Ontology Biological Processes (GOBPs) significantly enriched among the 35 upregulated differential proteins (DPs) identified in part (F) using the GOnet, with the 4,645 filtered proteins used as the background for the enrichment analysis. Details of proteins and GO terms in the network are included in Fig S1J.

Our data show a strong overlap between the biological replicates, with 4,645 proteins detected in at least three of the four experiments for either of the experimental conditions (i.e., Control or HSP90i) (Fig 1C, Figs S1A–B). For these 4,645 proteins (Table S4), there was a good pairwise correlation between the four replicates—both when comparing the summed Label-Free Quantitation (LFQ) intensities across all fractions, and LFQ intensities for each SEC fraction separately (Fig S1C). Using a heatmap to visualize scaled mean intensities for each of the 4,645 proteins across the 24 fractions (Fig 1D), we did not notice drastic differences between the Control and HSP90i condition—suggesting that HSP90 inhibition does not trigger widespread remodeling of the native proteome.

In order to discount the possibility that this lack of obvious changes to the SEC-proteome between our two conditions was due to lack of target modulation (i.e., HSP90 inhibition), we set out to confirm that tanespimycin treatment in our experiment led to the molecular changes expected in response to HSP90 inhibition. As our study is the first HSP90 inhibitor-based analysis of its kind, there were no other SEC-MS datasets for direct comparison. Therefore, we summed the individual intensities from all 24 fractions for the 4,645 filtered proteins, imputed missing values based on a left-shifted Gaussian distribution, and performed limma-based differential expression analysis using the *R* package DEP (Zhang et al., 2018) on these summed intensities (Fig S1D–H). In this way, we identified proteins whose abundances changed significantly between the Control and HSP90i conditions (Fig 1E–F, Fig S1I, and Table S5–S6)—in essence replicating previous bulk whole-proteome analyses of protein abundance changes in HSP90 inhibitor-treated cells (Fierro-Monti et al., 2013; Maloney et al., 2007; Quadroni et al., 2015; Savitski et al., 2018; Sharma et al., 2012; Schumacher et al., 2007; Voruganti et al., 2013; Wu et al., 2012). Of the 76 differential proteins (DPs) identified in our summed analysis (adjusted *p* < 0.05 and absolute log_2_(Fold Change) > 2) (Fig S1I), 47 DPs had also been identified in previous HSP90 inhibitor-proteomic studies (Fig 1F–G and Fig S1I, in green; Table S6), including 27 of the 41 down-regulated proteins. Furthermore, we confirmed induction of the heat shock response—triggered by activation of HSF1 following HSP90 inhibition (Bagatell et al., 2000)—as indicated by the presence of 20 HSF1 targets (Kovacs et al., 2019) among the 35 up-regulated proteins (Fig 1F–G magenta outlines, Fig S1I magenta stars, Table S6), and corresponding enriched Gene Ontology Biological Processes (Pomaznoy et al., 2018) such as ‘response to heat’, ‘response to unfolded protein’, and ‘chaperone mediated protein folding’ (Fig 1H, Fig S1J). Therefore, we were confident that we had achieved HSP90 inhibition in our experiment.

### Distinct SEC profiles and subtle changes following HSP90 inhibition for HSP90 machinery subunits

Having confirmed that the proteins whose overall summed abundances changed in our dataset were consistent with previous bulk proteomic studies of HSP90 inhibitor-treated cell lines, we proceeded with the aspect that was unique to our study: the fact that we had SEC traces for each individual protein. We first focused on the same two molecular hallmarks of HSP90 inhibitor treatment that we had evaluated with the summed differential protein analysis, i.e., increased levels of the HSF1 target gene products HSP72, HSP27, and BAG3, and depletion of the HSP90 client proteins CDK1, CDK6, and CHEK1, as well as AKT1 (Fig 2A–B, Fig S2A). In both cases, the SEC-MS profiles demonstrated clear and consistent trends in line with the summed analysis. Products of HSF1 target genes showed a general increase in all SEC fractions, as opposed to an increase in any particular sub-population (e.g., higher molecular weight oligomers that represent the active form of small heat shock proteins (Haslbeck et al., 2019))(Fig 2A, Fig S2A). Interestingly, we note that the profile of HSF1 itself did not differ appreciably between Control and HSP90i conditions (Fig S2A), despite canonical models of heat shock response induction involving HSF1’s dissociation from HSP70 and/or HSP90 prior to trimerization, activation, and nuclear translocation (Masser et al., 2020; Pincus, 2020). In contrast to the aforementioned proteins, HSP90 clients generally eluted in one clear peak near the estimated molecular weight for the monomeric species, which had lower intensities in the HSP90i condition—consistent with client protein destabilization and degradation (Fig 2B). We validated the global SEC-MS data by repeating our HSP90 inhibitor treatment and initial SEC-fractionation, followed by SDS-PAGE and immunoblotting for individual proteins (SEC-IB). In the majority of cases, our targeted immunoblot analysis correlated well with the mass spectrometry readout. For example, SEC-IB confirmed the increase across all fractions in HSP72 and BAG3, with the loading control GAPDH remaining unchanged (Fig 2A).

**Figure 2.**
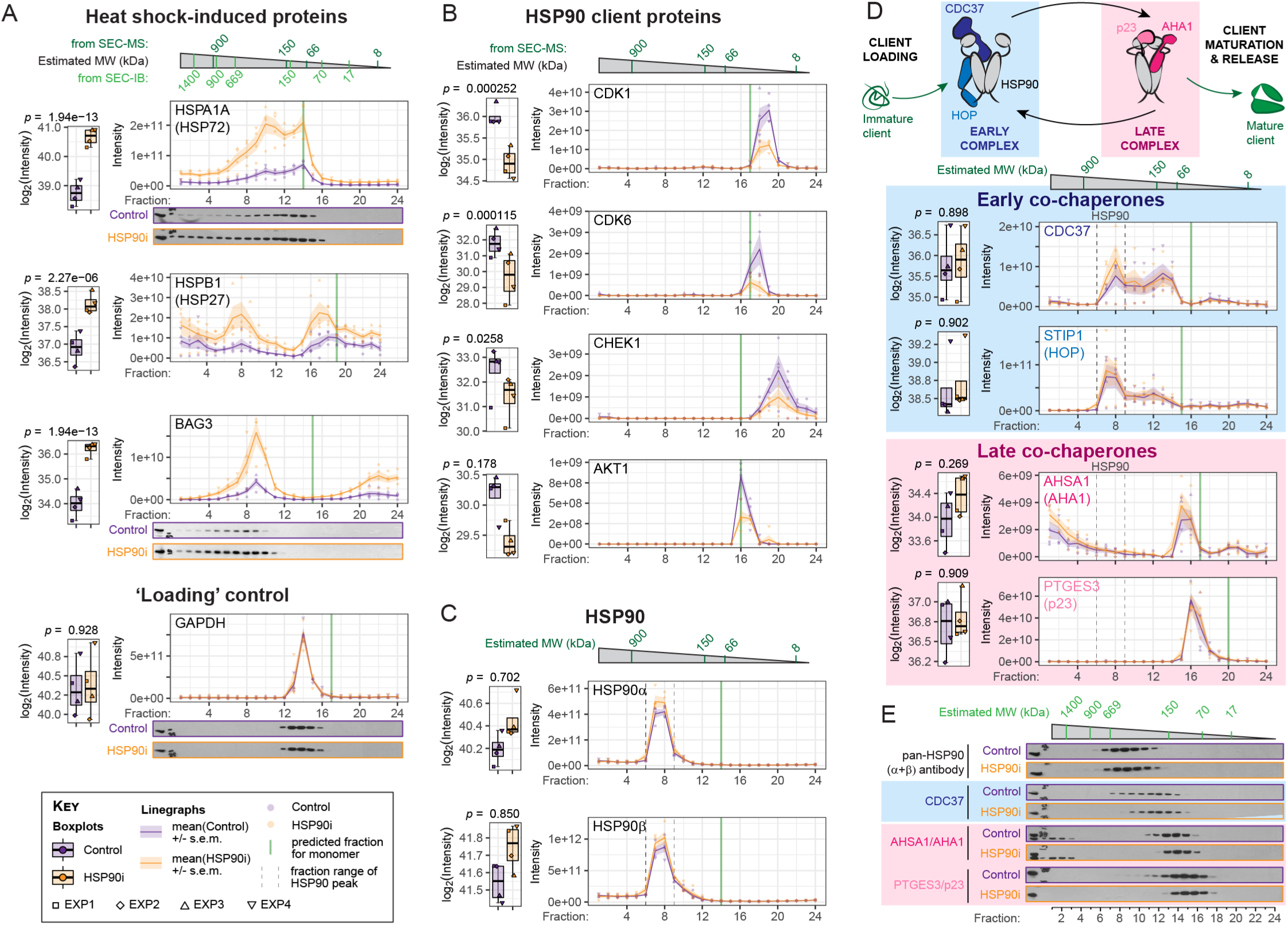
Profiling changes in the HSP90 machinery. (**A–B**) Tanespimycin-induced induction of heat shock factor-1 (HSF1) regulated proteins HSP70/HSP72 and BAG3 (A), and depletion of the HSP90 client proteins CDK1, CDK6, CHEK1, and AKT1 (B), observed in the SEC-MS dataset, and confirmed by SEC-Immunoblot (SEC-IB). GAPDH levels were used as a ‘loading’ control. (**C**) Both inducible (HSP90AA1/HSP90α) and constitutive (HSP90AB1/HSP90β) cytoplasmic HSP90 isoforms elute predominantly between fractions 6 and 9, and do not change significantly following HSP90 inhibitor treatment. (**D**) HSP90 co-chaperones considered to play a role early in HSP90’s ATP-dependent client maturation cycle (blue hues) co-elute with HSP90, whereas later co-chaperones (pink hues) do not. See also Fig S2. (**E**) SEC-IB profiles of the HSP90 machinery identifies subtle yet clear shifts of co-chaperones to lower molecular weight fractions. All SEC-Immunoblots are representative of three independent biological replicates.

Switching focus to HSP90, the two major cytoplasmic HSP90 isoforms were distributed almost exclusively in fractions 6–9 for both SEC-MS and SEC-IB readouts (Fig 2C)—consistent with a high molecular-weight oligomeric complex observed at 400– 500 kDa by targeted native complex separation approaches (Eskew et al., 2011; Moullintraffort et al., 2010). Note that these HSP90 profiles indicate that a negligible fraction of the total HSP90α and HSP90β population is present solely as dimers or monomers in HT29 cells, consistent with higher-order HSP90-containing ‘epichaperome’ assemblies in cancer (Kamal et al., 2003; Rodina et al., 2016). Note also that the ER-resident HSP90 isoform HSPB1/GRP94 was also present almost exclusively in the higher molecular weight fraction range, whereas the mitochondrial HSP90 TRAP1 mostly co-eluted as a monomer (Fig S2B)— a finding that was also apparent when we plotted the SEC-MS profiles of these HSP90 isoforms in a HeLa-CCL2 dataset (Heusel et al., 2020)(Fig S2C). By contrast, the profiles of HSP72 and HSP27 had multiple peaks representing both monomeric and higher molecular weight populations (Fig 2A). Also in contrast with the other heat shock proteins, neither the profiles nor total abundance of the HSP90s changed between control and treated conditions (Fig 2C, Fig S2B)—a finding that is consistent with previous observations that HSP90 inhibition does not induce HSP90 expression (Maloney et al., 2007; Ghalhar et al., 2014; Karkoulis et al., 2010; Miao et al., 2019).

HSP90 functions as part of a large multi-protein complex, consisting of dozens of co-chaperones that are required for various parts of its ATP-dependent client activation cycle (Schopf et al., 2017). We had previously shown that the HSP90 cochaperones HOP, CDC37, AHA1, and p23 all dissociate from HSP90-client protein complexes in HT29 cells under the same tanespimycin treatment conditions employed in the present study (Samant et al., 2014). Therefore, we were surprised to find minimal changes to the SEC profiles of HSP90 co-chaperones following HSP90 inhibition (Fig 2D, Fig S2D–E). Profiles of the co-chaperones could be grouped according to their role in HSP90’s ATP-driven client activation cycle. The early co-chaperones STIP1/HOP and CDC37, both involved in the loading of client proteins onto the chaperone, had major peaks that overlapped with HSP90 (i.e., fractions 6–9)(Fig 2D). Later co-chaperones that are involved in client maturation and release (e.g., AHA1, p23) did not co-fractionate with HSP90 and were found in lower molecular weight fractions (Fig 2D, Fig S2D). Note that the protein phosphatase PPP5C/Ppt1, which de-phosphorylates both HSP90 and CDC37 prior to ATP hydrolysis by the chaperone machinery (Soroka et al., 2012; Vaughan et al., 2008), also coeluted with the HSP90α/β peak (Fig S2D). Our findings suggest that interactions between HSP90 and the late co-chaperones are not preserved through the sample preparation protocol, and perhaps that these interactions are weaker or more labile than those between HSP90 and the early co-chaperones. Interestingly, of the two main E3 ubiquitin ligases associated with the HSP90 machinery, the tetratricopeptide (TPR) motif-containing co-chaperone CHIP/STUB1 co-fractionated with HSP90, whereas CUL5 did not (Fig S2E).

Following up the SEC-MS with SEC-IB analysis, we noticed subtle yet clear changes to the distribution of co-chaperones—both early and late—following HSP90 inhibition, with the fraction distributions becoming narrower in the treated samples (Fig 2E). For both early (CDC37) and late (AHA1, p23) co-chaperones, the tightening of the distribution was skewed towards the lower molecular weight fractions, suggesting that the higher molecular weight fractions—which overlapped with the lower end of the HSP90 SEC-IB distribution—were disrupted following HSP90 inhibition, in line with our previous observations (Samant et al., 2014). This interpretation would also suggest either that most of the co-chaperone population in both control and treated cells is not in complex with HSP90, or that the majority of HSP90 : co-chaperone interactions are too labile to withstand sonication and fractionation, as required for the SEC protocol.

### Identification of differential protein complexes based on SEC coelution feature detection

The observation that certain HSP90 complex subunits co-eluted, whereas others did not, drove us to determine more globally the degree to which protein complexes were preserved in our dataset. According to the *R* package *CCprofiler* —developed to analyse SEC-MS data (Heusel et al., 2019)—an estimated 39–46 % of the protein mass was in the ‘assembled’ (*vs*. monomer) fraction size range in our complete dataset of 6,427 proteins, with good consistency across all eight samples (Fig 3A). The dataset contained 1,796 of the 2,532 proteins annotated in the CORUM protein complex database (Ruepp et al., 2010)(Fig S3A), with 1,457 of the 1,753 CORUM-annotated protein complexes represented at 50 % subunit coverage or more (Fig S3B). When assessing coeluting subunits from the SEC fraction profiles, *CCprofiler* identified 247 CORUM protein complexes, defined as at least 50 % of the CORUM annotated subunits classified as co-eluting at 5 % FDR (Fig 3B). An alternative SEC-MS protein complex predictor, PCprophet (Fossati et al., 2021), identified 408 CORUM complexes with the same threshold of at least 50 % of subunits co-eluting (Fig 3B).

**Figure 3.**
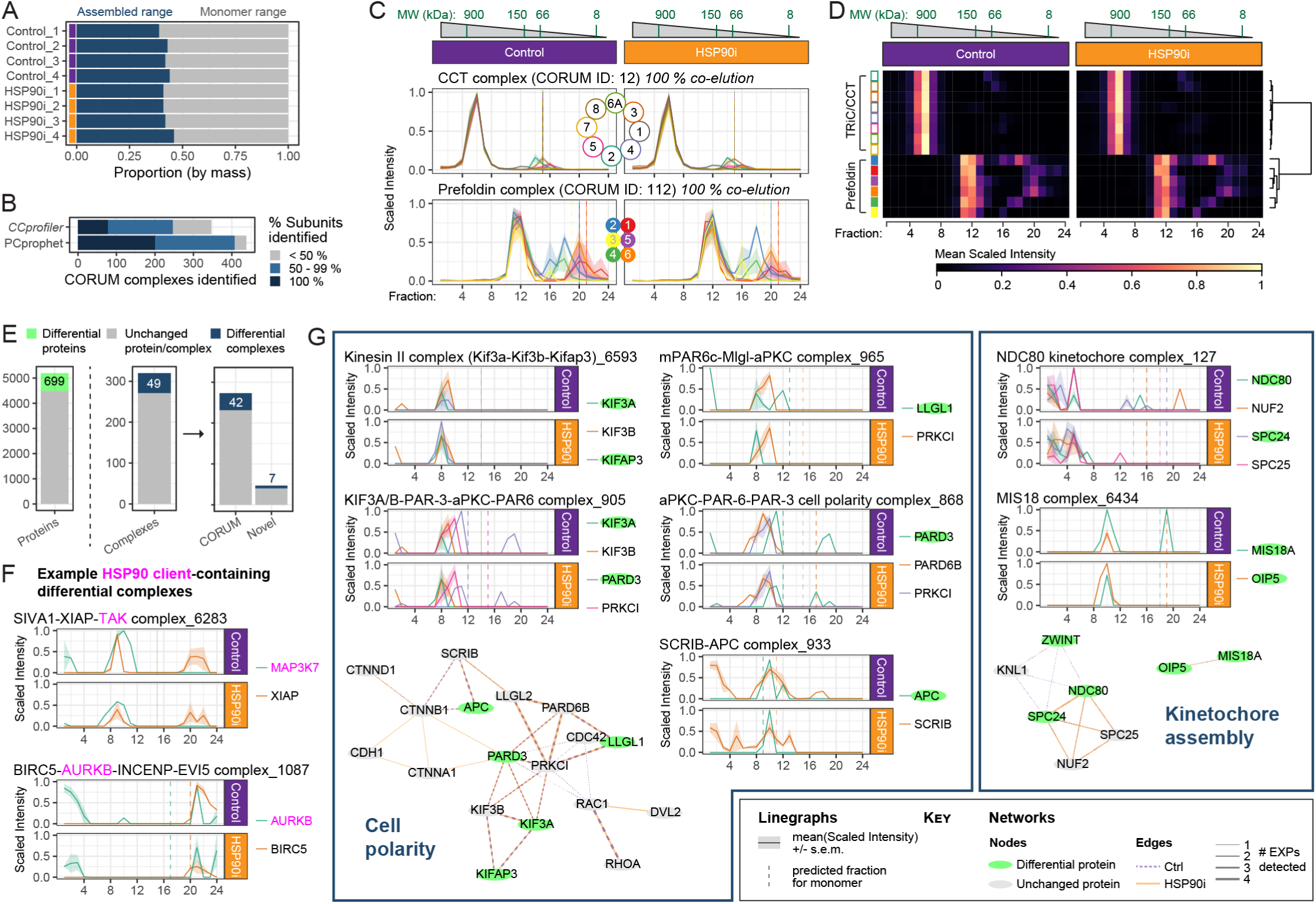
Complex-centric analysis to identify CORUM protein complexes in SEC-MS dataset. **(A)** Global statistics of the proportion of the protein signal in each sample attributed to assembled or monomeric state in our dataset of 6,427 proteins, as estimated by *CCprofiler*. **(B)** Number of CORUM-annotated protein complexes identified based on co-elution of the SEC-MS profiles of their constituent subunits, using *CCprofiler* or PCprophet packages. The percentage of subunits identified are also indicated. (**C**) Mean scaled intensity profiles for each subunit of the hetero-oligomeric chaperones TRiC/CCT (top) and Prefoldin (bottom). Both complexes were fully detected by *CCprofiler*, but the holo-enzyme consisting of Prefoldin and TRiC/CCT was not detected. (**D**) Heatmap of mean scaled intensities for TRiC/CCT and Prefoldin complex subunits. The dendogram cut-offs based on Euclidean distance matrix with the Ward-D2 linkage method are illustrated to the right of the heat map. Similar data for the 26S proteasome is shown in Fig S3C. (**E**) Number of proteins (left) and protein complexes (right) identified by PCprophet as being differentially regulated following HSP90 inhibition. The protein complexes identified were further separated as those annotated on CORUM, and those that were ‘novel’ complexes. See Table S7 & S8 for full list of proteins and protein complexes, respectively, identified by PCprophet. (**F**) Mean scaled intensity profiles for each identified subunit of the differential protein complexes SIVA1–XIAP–TAK (CORUM ID: 6283) and BIRC5–AURKB–INCENP–EVI5 (CORUM ID: 1087). Established HSP90 clients are indicated in magenta. (**G**) Mean scaled intensity profiles (from Table S3) for selected differential protein complexes, grouped according to their biological function. Protein–protein interaction networks consisting of subunits within these complexes are also shown. Green nodes represent differential proteins identified using PCprophet’s protein-centric analysis. Edge width represents the number of experiments in which the interaction was confidently detected by PCprophet, with the edge colour representing Control or HSP90i detections. Profiles of all 49 differential complexes are shown in Fig S4, and all protein–protein interactions networks identified by PCprophet in Fig S5. Dashed vertical lines on linegraphs indicate the fraction in which the monomer would be detected, based on the UniProt-annotated molecular weight.

The discrepancy between protein complex coverage and co-elution in our dataset—and, indeed, in all SEC-MS studies to date (Fossati et al., 2021; Heusel et al., 2019, 2020; Kirkwood et al., 2013; Larance et al., 2013, 2016)—is likely due to a number of factors. For one, not all CORUM-annotated complexes will be present in all cell types. More importantly, the lysis conditions used— involving considerable sample processing time prior to SEC-based separation—will not preserve highly dynamic, transient, or labile interactions. To illustrate the second point, the hetero-oligomeric chaperones TRiC/CCT and Prefoldin co-eluted fully as individual protein complexes (major peaks in fractions 4–8 and 10–14, respectively), but the higher-order holoenzyme CCT-Prefoldin (Gestaut et al., 2019) was not preserved (Fig 3C–D). Similarly, the majority of 20S proteasome core particle subunits eluted as a peak distinct from the 19S regulatory particle (Fig S3C), suggesting that the 26S proteasome holoenzyme (Bard et al., 2018) is disrupted during sample processing. The 245 complexes *CCprofiler* identified in our dataset is of a similar order of magnitude as other SEC-MS studies (Fig S3D), although the higher numbers in those studies were presumably as a result of smoother elution profiles by SEC into more fractions, and/or fewer missing values due to use of data-independent acquisition for mass spectrometry (DIA-MS). Indeed, when Heusel *et al*. re-analysed their SEC-fractionated HEK293 cell lysate with data-dependent acquisition mass spectrometry (DDA-MS) instead of DIA-MS, they observed a > 50 % drop-off in CORUM complex identification (298 *vs*. 621) using the same *CCprofiler* parameters (Heusel et al., 2019)(Fig S3D).

Using PCprophet’s differential analysis workflow, we identified 699 proteins and 49 protein complexes as being significantly altered between our two conditions (Fig 3E, Tables S7–S8). As a percentage of the total positive IDs in the analysis, both the altered proteins (13.4 %) and complexes (15.3 %) were lower than that observed by Fossati *et al*. using the same PCprophet analysis workflow for comparing HeLa-CCL2 cells at interphase *vs*. mitosis (approximately 30 % and 26 %, respectively)(Fossati et al., 2021). This was not necessarily surprising, given that HSP90 is not a typical molecular chaperone responsible for general folding of the majority of the proteome (as opposed to HSC70, for example), but rather has a small subset of clients. Therefore, the changes induced by HSP90 inhibition would not be as widespread as the difference between two cell-cycle states.

The 49 HSP90i-modulated protein complexes identified through PCprophet spanned diverse biological processes, consistent with the diverse functional nature of HSP90’s clientele (Fig S4). Several of these protein complexes contained *bona fide* HSP90 clients, e.g., the oncoprotein kinases TAK1/MAP3K7 (Liu et al., 2008; Shi et al., 2009), Aurora kinase B (Lange et al., 2002; Terasawa and Minami, 2005), and class 3 PI3-kinases PIK3C3 and PIK3R4 (Giulino-Roth et al., 2017; Moulick et al., 2011), as well as their associated tumor suppressor Beclin1/BECN1 (Hasan et al., 2020; Xu et al., 2011)(Fig 3F, Fig S4). Furthermore, multiple signaling hubs known to require HSP90 activity were represented, including several protein complexes central to cell polarity (Baas et al., 2004; Benitez et al., 2014; Berezuk and Schroer, 2004) and kinetochore positioning during mitosis (Niikura et al., 2006)(Fig 3G). Importantly, not all of these HSP90i-modulated complexes are known direct HSP90 interactors. For example, neither of the HSP90i-modulated kinetochore-related complexes identified—MIS18 and NDC80—contain known HSP90 client proteins. Rather, HSP90 inhibition is proposed to impair kinetochore formation via destabilization and degradation of the MIS12 complex subunit DSN1 and/or the centromere-localised Polo-like kinase PLK1 (Davies and Kaplan, 2010; McKinley and Cheeseman, 2014; Niikura et al., 2017)—neither of which belong to NDC80 or MIS18 complexes. The identification of such indirect HSP90-dependent complexes highlights the utility of our SEC-MS approach, as they would not have been identified using HSP90interactomics, nor by bulk differential proteomics (as the total abundance of the constituent subunits did not change significantly upon HSP90 inhibition).

### Distinct HSP90 inhibitor-induced protein hits identified by summed *vs*. individual fraction intensity-based differential analysis

Despite the identification of multiple protein complexes known to require HSP90 for their function (both directly and indirectly) using PCprophet’s differential analysis workflow, we were concerned that the limited number of total protein complexes positively identified would result in us missing important protein– protein interaction changes for proteins not assigned to a specific complex. Therefore, we complemented the automated PCprophet analysis by searching for differences between Control and HSP90i-treated samples at the SEC fraction level using the same differential analysis workflow we had used for the summed intensities in Fig 1E–H, i.e., treating our dataset as 24 different Control *vs*. HSP90i comparisons (Fig 4A).

**Figure 4.**
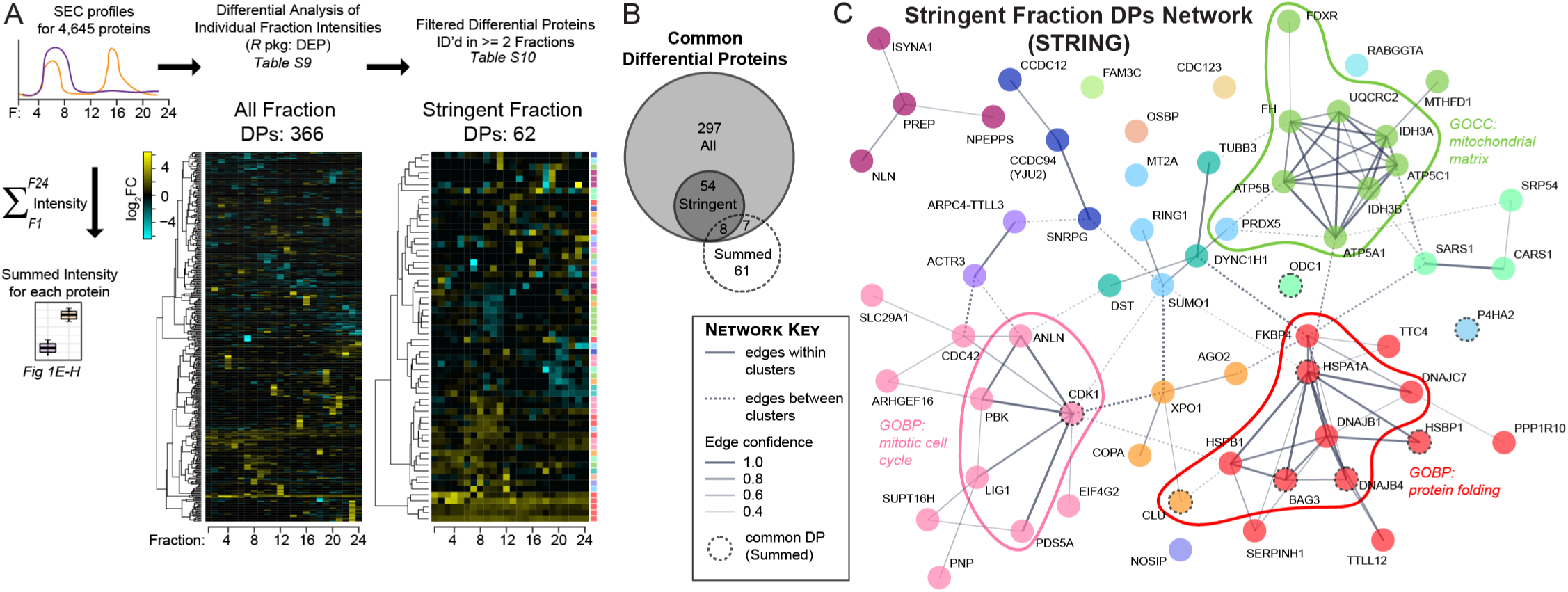
Identifying proteins whose SEC profiles changed following HSP90 inhibition at an individual fraction level. (**A**) Approach to calculate differential proteins based on individual SEC fraction intensities between Control and HSP90i conditions, using the *R* package DEP. Heatmaps depict log_2_ FC for the 366 proteins identified as hits (absolute log_2_ FC > 1 and adjusted *p*-value < 0.05) in any of the 24 SEC fractions (‘All Fraction DPs’, left), and those identified as DPs in two or more fractions (‘Stringent Fraction DPs’, right), using the *R* package DEP. Hits were clustered (Euclidean distance with Ward-D2 linkage) before plotting as heatmaps. Colours to the right of the Stringent Fraction DPs heatmap represent Markov clusters calculated on STRING-db with an inflation parameter of 2 (see also part (C)). (**B**) Venn/Euler diagram (*R* package ‘eulerr’) representing the number of common proteins between the 366 All Fraction DPs and 62 Stringent Fraction DPs in (A), and 76 summed intensity-based DPs from Fig 1. (**C**) Network generated by STRING protein–protein interaction database using the 62 Stringent Fraction DPs as the input. Colours represent clusters from Markov clustering (inflation parameter = 2). Solid and dashed lines indicate protein–protein interactions (edges) within and between clusters, respectively; weight of lines between nodes indicate strength of evidence for the interaction on STRING (both functional and physical protein associations). Proteins also identified via summed intensity analysis are depicted with dashed circular outlines.

We identified 366 unique DPs across all 24 fractions (Fig 4A, Table S9). The majority of these proteins were only identified as DPs in one of the 24 fractions. Although single fraction changes could be indicative of important biological perturbations, we decided to filter out such singleton DPs in order to increase stringency and minimise the effect of stochastic fluctuations during SEC fractionation. After this step, we were left with 62 proteins that were DPs in two or more fractions (Fig 4A, Table S10).

We were surprised to see only eight common proteins between the 62 stringent fraction DPs and the 76 summed DPs from Fig. 1E–H (Fig 4B). Of these common proteins, all except one—CDK1, an HSP90 client—are known HSF1 target genes (Kovacs et al., 2019). Indeed, the only enriched GOBP term in the set of 62 stringent fraction DPs was ‘chaperone-mediated protein folding’ (adjusted *p* = 7.53e-7). This approach provides further evidence that, with the exception of the most abundant heat shock proteins, our SEC-based approach identifies a distinct set of proteins compared to those that would be identified using total cell lysate-based comparative proteomics.

Next, we generated protein–protein interaction networks for the 62 stringent DPs using either the STRING protein–protein interaction database (Szklarczyk et al., 2021) or the curated interactome in the canSAR database (Mitsopoulos et al., 2021) (Fig 4C, S6). Both tools yielded networks with significantly more edges than expected by chance, i.e., 47 edges expected by chance; 99 edges between 54 nodes observed with STRING (*p* = 3.25e-11); 76 edges between 43 nodes observed with canSAR. These highly-connected networks suggested that the 62 DPs as a group were biologically related. STRING’s built-in Markov Cluster Algorithm (Enright et al., 2002) identified three large clusters (those with > five nodes each); the largest of these clusters (Fig 4C, red nodes) was heavily enriched in molecular chaperones and co-chaperones (10 of the 12 cluster nodes), and included four of the eight DPs common with the summed DPs (Fig 4C, dashed node outlines). Another cluster contained numerous cell cycle and cytoskeletal proteins (Fig 4C, pink nodes). The final large cluster consisted of nine mitochondrial proteins (eight from the mitochondrial matrix), including members of the ATP5 synthase complex (3/6 subunits) and isocitrate dehydrogenase 3 (IDH3) complex (2/3 subunits)(Fig 4C, light green nodes).

### The actin-binding oncoprotein Anillin is recruited to inhibited HSP90 complexes

As the largest cluster from our Stringent Fraction DP network (Fig 4C, red nodes) consisted exclusively of well-characterised heat shock response proteins, we decided to focus on the other two clusters to gain potentially novel biological insight. Of the four highest-confidence interactions within the cell cycle and cytoskeletal protein cluster (STRING edge confidence > 0.9), CDK1, PBK, and PDS5A were previously identified as HSP90 inhibitor-modulated proteins (Sui et al., 2020; Wu et al., 2012). We therefore probed further into the fourth node in this high-confidence edge network: ANLN/Anillin, a cytoskeletal scaffold protein that links RhoA, actin, and myosin during cytokinesis.

Anillin is emerging as an important regulator of the Epithelial-to-Mesenchymal Transition (EMT) during malignancy (Naydenov et al., 2021; Xu et al., 2019), and is upregulated in a variety of tumors—including colon cancer (Lian et al., 2018; Wang et al., 2016; Zhang et al., 2018a,b). Here, Anillin almost met our hit criteria in the initial global analysis of summed intensity changes (Fig 1E–H), missing out because it fell just under the fold change threshold (Fig 5A boxplot, *p* = 0.0117, log_2_FC = 0.86). The degree of Anillin modulation was strikingly consistent with the HSP90i-induced Anillin changes observed when we analysed previous proteomics datasets in HeLa cervical (*p* = 0.0002, log_2_FC = 0.94), MDA-MB-231 triple-negative breast (*p* = 0.0411, log_2_FC = 0.92), and CAL-27 oral squamous (*p* = 0.0007, log_2_FC = 1.28) carcinoma cell lines (Sharma et al., 2012; Wu et al., 2012). Using our individual fraction-based approach, Anillin was a hit in fractions 6 and 7 (Fig 5A linegraph, asterisks). Although Anillin’s SEC profile at first glance looked typical of HSF1-induced proteins, we noticed a subtle shift in the major peak towards higher molecular weight fractions in the HSP90i conditions (Fig 5A, linegraph), as opposed to a global increase across almost all fractions (e.g., compared with HSP72, HSP27, and BAG3 SEC profiles in Fig 2A). This shift was even more apparent with an SEC-IB readout (Fig 5A, immunoblots). As this shift resulted in more of the Anillin population overlapping with the HSP90 peak (Fig 5A, linegraph, dashed lines), it occurred to us that Anillin might be interacting with HSP90 complexes, and that this interaction was increased upon HSP90 inhibition. Indeed, immunoprecipitation of endogenous HSP90 confirmed that Anillin is recruited to HSP90-containing complexes following tanespimycin treatment (Fig 5B).

**Figure 5.**
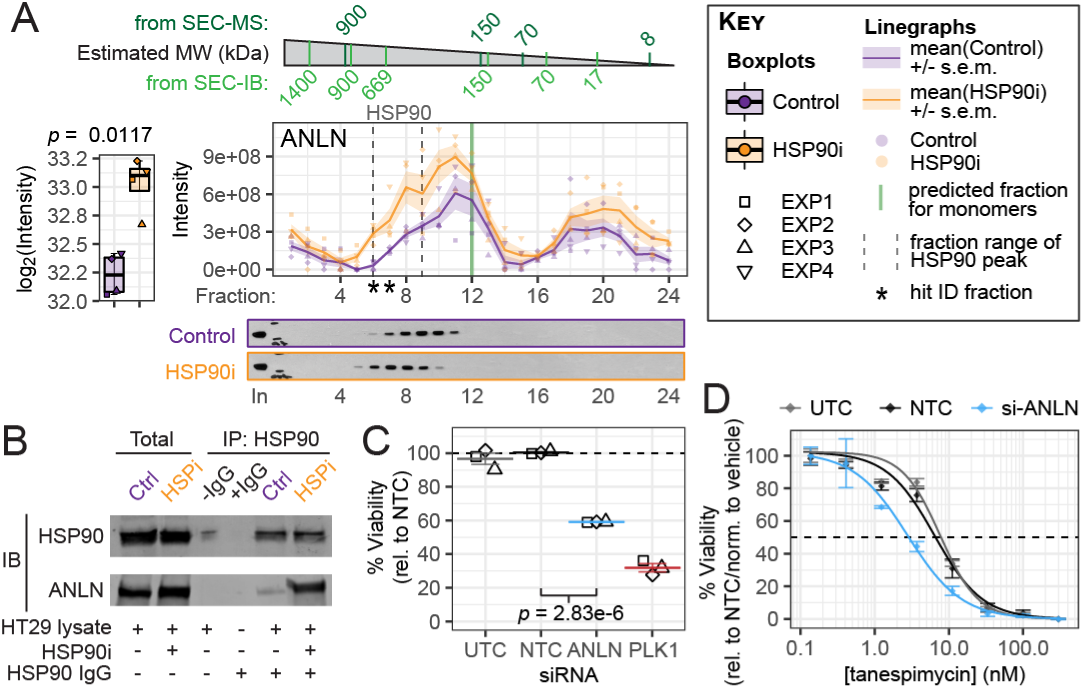
Actin-binding protein Anillin/ANLN is recruited to inhibited HSP90 complexes. **(A)** Left, total ANLN/Anillin protein levels increase following HSP90 tanespimycin treatment (*p* = 0.0017, log_2_ FC = 0.86). Right, SEC-MS and SEC-IB elution profiles indicate a shift towards higher molecular weight complexes for the major Anillin peak following HSP90 inhibition. This includes an increase in the same fractions as the HSP90 elution peak (linegraph, dashed lines). (**B**) Anillin co-immunoprecipition with HSP90 is increased following tanespimycin treatment. HT29 human colon cancer cells were treated with 5 x GI_50_ (62.5 nM) tanespimycin, or mock-treated with the same volume of vehicle DMSO control, for 8 h. HSP90 immunoprecipitation (IP) was performed on 1 mg of lysate protein from each condition, and the resultant immunoblots (IB) were probed for HSP90 and Anillin, as indicated. 15 µg of lysate protein was loaded for the ‘Total’ lanes. (**C, D**) Knockdown of Anillin reduces HT29 cell viability and results in a two-fold sensitization to tanespimycin. HT29 cells were treated for 48 h with siTOOLs pool of 30 siRNAs (25 nM total concentration) targeted against Anillin (ANLN), non-targeting control NTC, untargeted control UTC, or death-inducing control PLK1, followed by treatment for 96 h with a range of tanespimycin concentrations (except for PLK1), or mock-treatment with vehicle control (0.1 % DMSO), while still in the presence of the original siRNAs. Confluency was monitored every 8 h throughout the time-course by Incucyte® (see Fig S7). Cell viability at the end of the time-course (144 h total) was measured using CellTiter-Blue® cell viability assay. Cell viability results for the mock vehicle-treated controls only are plotted in (**C**), with adjusted *p*-value (as determined by one-way ANOVA with Tukey’s multiple comparison test) for Anillin *vs*. NTC. Lines and error bars represent mean +/- standard error for each condition from three biological replicates (EXP1–EXP3, different shapes). Plotting all the dose-response data (**D**) identified two-fold sensitization to tanespimycin treatment on Anillin knockdown in HT29 cells, with GI_50_ reduced to 2.9 nM (*vs*. 6.6 nM for NTC). Points and error bars represent mean +/- standard error for each condition. Dose-response curve fitting was performed using the ‘Log[Inhibitor] *vs*. normalized response – Variable slope’ non-linear regression model in Graphpad Prism. See Fig S7C for dose-response curves relative to vehicle-treated (0.1 % DMSO) NTC control.

We hypothesized that recruitment of Anillin to inhibited HSP90 complexes may be an important part of the cellular response to HSP90 inhibition. Knockdown of Anillin by pooled siRNAs for six days resulted in a 42 % and 49 % reduction in HT29 cell viability and confluency, respectively, *vs*. the negative non-targeting control (Fig 5C, S7A–B). Although this reduction in cell viability made it difficult to assess any further effect of Anillin knockdown on HSP90 inhibitor sensitivity (Fig S7C), we identified a two-fold reduction in tanespimycin GI_50_ concentration to 2.9 nm (*vs*. 6.6 nm for the negative non-targeting control siRNAs)(Fig 5D). These findings lend further weight to the role of Anillin in malignancy (Naydenov et al., 2021; Tuan and Lee, 2020), and suggest effects on HSP90-mediated networks as at least a partial mechanism for its pro-tumor function.

### Mitochondrial IDH3 is a novel HSP90-dependent protein complex

Finally, switching focus to the mitochondrial matrix-enriched cluster from our Stringent DP Network (Fig 4C), we were especially interested in the IDH3 complex, as the third subunit was identified in our less stringent list of 366 DPs at the individual fraction level. The total levels of any of the three subunits of the IDH3 complex did not change (*p* > 0.05 and absolute log_2_FC < 1) (Fig 6A, boxplots). Therefore, the IDH3 complex would not be identified as a hit through traditional bulk proteomics. Plotting the SEC-MS traces, however, brought to light clear changes in a certain sub-population of IDH3 subunits in HSP90i-treated cells (Fig 6A, linegraphs). The bulk of the signal for all three subunits was in low molecular weight fractions, most likely representing the monomeric species. These peaks were the same in control and HSP90 inhibitor-treated conditions. However, a minor peak around fraction 10—representing a larger molecular weight complex of approximately 300 kDa—was clearly reduced following HSP90 inhibition for all three subunits. This would be consistent with the size of an octameric complex, thought to represent the active form of IDH3 in cells (Huh et al., 1997).

**Figure 6.**
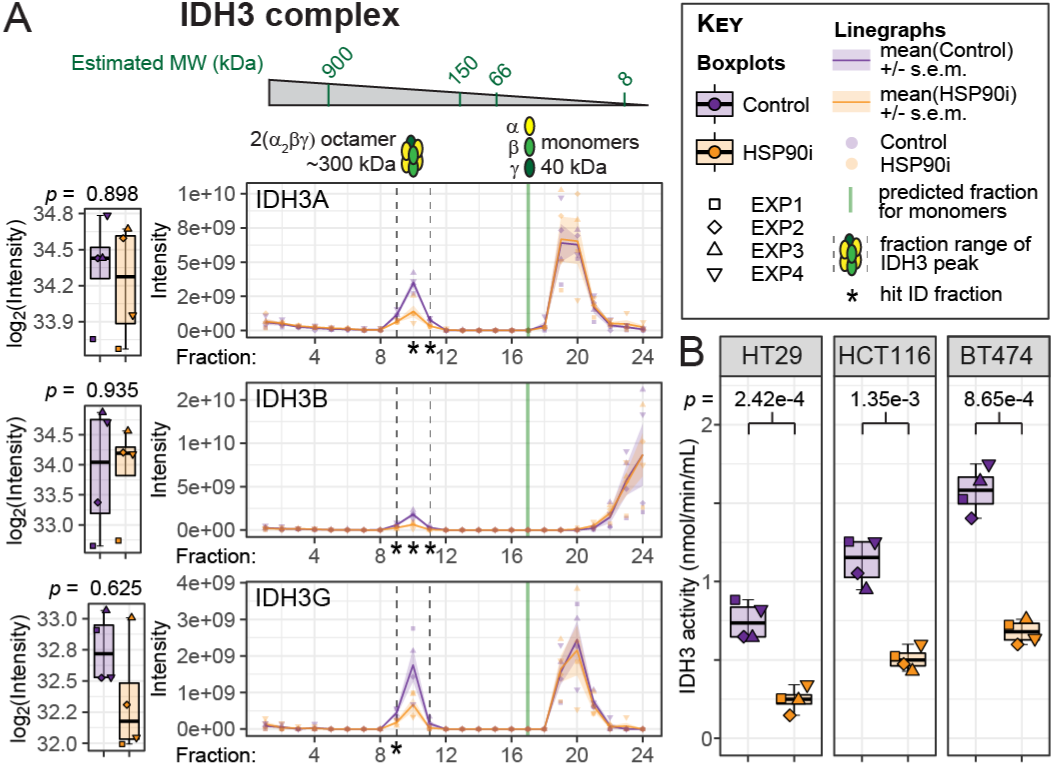
Mitochondrial isocitrate dehydrogenase 3 (IDH3) complex is disrupted upon HSP90 inhibition. **(A)** Left, Box/Tukey plots illustrate that total levels of the three IDH3 protein complex subunits do not change following HSP90 inhibition. Right, SEC profiles identify a high molecular weight peak (corresponding to the size of the octameric IDH3 complex) that is significantly reduced for all three subunits in HSP90 inhibitor-treated cells. Asterisks depict significantly differential fractions. (**B**) IDH3 activity is significantly reduced upon HSP90 inhibition in HT29 human colon adenocarcinoma, HCT116 human colon carcinoma, and BT474 human breast ductal carcinoma cell lines. Activity was measured using the IDH Activity Assay Kit (Sigma) according to manufacturer’s instructions, using NAD+ as the co-factor. See Fig S8A for the corresponding assay with NADP+ as the co-factor, for estimating IDH1 and IDH2 activity. Adjusted *p*-values (two-tailed Student’s *t* -test) are shown.

We reasoned that HSP90 could play an important role in the maintenance of active IDH3 complexes. We showed using a cell-based reporter assay that IDH3 activity is reduced by >50 % following HSP90 inhibition in HT29 cells, as well as in two other cell lines—the HCT116 colon carcinoma, and BT474 breast ductal carcinoma (Fig 6B). The effect was specific to this IDH complex, as the activities of IDH1 and IDH2—which use NADP^+^ as a co-factor, rather than NAD^+^––were unaffected (Fig S8A).

Finally, we confirmed that this was a general effect of HSP90 inhibition by performing the same assay in cells treated with two other HSP90 inhibitor chemotypes in HCT116 cells (Fig S8B). We therefore identify the IDH3 complex as a novel component of the HSP90-dependent proteome that would most likely have remained uncharacterised using traditional bulk proteomics or interactomics approaches.

## Discussion

Defining the scope of the HSP90-dependent proteome has been a subject of enquiry in both basic and translational biology for over a decade (Citri et al., 2006; Hartson and Matts, 2012; Maloney et al., 2007; Samant et al., 2012; Schumacher et al., 2007; Weidenauer et al., 2017). Here, we present an alternative approach to expand insight into this challenge, employing changes in the native complex profiles of the proteome rather than relying on changes in bulk abundance or direct interacting partners of HSP90. After confirming target engagement (i.e., HSP90 inhibition) in our dataset through summed differential protein analysis (Fig 1, Fig S1), we discovered through plotting SEC profiles of the HSP90 machinery that certain interactions were preserved, whereas others were almost entirely lost in these experiments (e.g., HSP90 interactions with early *vs*. late co-chaperones) (Fig 2, Fig S2). This ‘all-or-nothing’ phenomenon also held true at a global level: numerous obligate hetero-oligomeric protein complexes (e.g., TRiC/CCT and Prefoldin; 20S and 19S proteasome) co-eluted, whereas larger holo-complexes (e.g., CCT-Prefoldin; 26S proteasome) did not (Fig 3C–D, Fig S3C). Employing a complex-centric peak co-elution approach through the packages *CCprofiler* and PCprophet, we identified 49 HSP90i-modulated protein complexes between the Control and HSP90i condition (Fig 3E–G, Fig S4). These included several protein complexes containing known HSP90 clients, as well as protein complexes involved in downstream HSP90-dependent biological processes. To complement this complex-centric approach, we performed differential expression analysis at an individual fraction level, yielding 62 stringent differential proteins (Fig 4). Finally, we validated two novel hits from these stringent fraction-level differential proteins—Anillin (Fig 5, Fig S7) and the IDH3 complex (Fig 6, Fig S8)—identifying these as potentially important components of the HSP90-dependent proteome.

Our approach provides several novel insights into the HSP90 inhibition response that have so far remained uncharacterised through traditional comparative proteomics (e.g., mass spectrometry of Control *vs*. Treated whole cell lysates) or interactomics (e.g., mass spectrometry of HSP90 or co-chaperone coimmunoprecipitates; LUMIER and other bait : prey interaction screens). This was perhaps best exemplified by the limited overlap between differential proteins (DPs) identified with our individual fraction approach and those identified for summed intensities (Fig 4 *vs*. Fig 1E–H)—especially as the same workflow was used in both analyses. We demonstrate the utility of our dataset by identifying two as yet uncharacterized components of the HSP90-dependent proteome, that were only identified as DPs with our individual fraction-level analysis.

We discovered that Anillin abundance was significantly altered in previous HSP90 inhibition datasets (Sharma et al., 2012; Wu et al., 2012); yet it had not, to our knowledge, been validated in these or any other HSP90-related studies. We followed up this DP for two main reasons. In the context of the present study, Anillin’s shift to higher molecular weight fractions upon HSP90 inhibition was a relatively rare trait in the dataset. We therefore reasoned that Anillin might be recruited to specific complexes upon HSP90 inhibition—and even that this recruitment might occur at the level of HSP90 complexes directly, given the significant increase by fraction-level differential analysis in the HSP90 elution peak (Fig 5A). This was confirmed by co-immunoprecipitation of Anillin with HSP90, which increased following tanespimycin treatment (Fig 5B). The second reason for our interest in Anillin was a growing body of research showing its role in tumor progression, across a range of indications (Hall et al., 2005; Lian et al., 2018; Naydenov et al., 2021; Tuan and Lee, 2020; Wang et al., 2016; Xu et al., 2019; Zhang et al., 2018a,b). Our finding that Anillin knockdown for six days results in a smaller number of viable cells (Fig 5C) is consistent with this protein’s central role in cytokinesis (Piekny and Maddox, 2010). Although the effect of Anillin knockdown on sensitivity to tanespimycin was relatively modest (Fig 5D), it is suggestive of a potential relationship between HSP90 inhibitor efficacy and Anillin levels—which, as we have discussed, is overexpressed in several cancers. Anillin provides yet another potential link between the HSP90 machinery and cancer cell invasion, together with the cell polarity and kinetochore assembly-associated protein complexes identified as HSP90i-modulated by PCprophet in this study (Fig 3G).

Whereas careful mining of previous HSP90 inhibition proteomics datasets might have identified Anillin as a HSP90-modulated protein, the mitochondrial IDH3 complex has not been associated with the HSP90 inhibition response—and likely would have remained uncharacterised in this context through targeted interaction or differential abundance approaches. Importantly, we found that HSP90 inhibition specifically impaired activity of the IDH3 complex and not of the other two IDH family members IDH1 and IDH2 (Fig 6B, Fig S8). IDH3 is not as well characterized as IDH1 and IDH2 in the context of cancer and other diseases (Tommasini-Ghelfi et al., 2019); however, more recent studies implicate aberrant expression of IDH3—especially the alpha subunit—in malignancy (May et al., 2019; Zeng et al., 2015; Zhang et al., 2015). This is likely related to the fact that rewiring of energy metabolism is an extended hallmark of cancer (Hanahan and Weinberg, 2011). Indeed, the Warburg effect—where cells switch from mitochondrial oxidative phosphorylation to glycolysis as their predominant means of ATP production, even under oxygen-rich conditions—is commonly observed during the malignant process. As part of the tricarboxylic acid cycle, IDH3 would play a key role only in ATP production in cells that have not undergone Warburg-like metabolic transformations. Future studies could test whether the degree of glycolytic shift of a tumor correlates negatively with its sensitivity to HSP90 inhibitors. Furthermore, the fact that healthy cells are presumably more dependent on the tricarboxylic acid cycle—and therefore IDH3—than cancer cells might contribute to a narrowing of the therapeutic window for HSP90 inhibitors *in vivo*. More recent data suggest that IDH3 levels decrease during senescence (Cao et al., 2019)—a finding of potential importance in light of the interest in HSP90 inhibitors for targeting senescent cells (Fuhrmann-Stroissnigg et al., 2018).

Despite the novel biology revealed in this study, several limitations of SEC-MS—as well as potential improvements—became apparent during our analysis. Almost 1,500 CORUM protein complexes were detected at greater than 50 % subunit coverage (Fig S3B), yet *CCprofiler* and PCprophet assigned only 245 and 408 CORUM complexes, respectively, as having co-eluting SEC traces (Fig 3B). This number was slightly lower than existing published SEC-MS studies (Fig S3E), which could be caused by a number of technical and/or biological differences in our dataset. For example, at the biological level, HT29 colon carcinoma cells could have fewer or more labile protein complexes than HEK293 and HeLa cells—two workhorse cell lines with wide-ranging abnormalities at the genetic and protein level (Landry et al., 2013; Lin et al., 2014). At the technical level, a key component of unbiased complex-centric proteome profiling algorithms such as those implemented by *CCprofiler* and PCprophet is the consistent identification of the same set of target proteins in consecutive SEC fractions (Heusel et al., 2019). Co-elution feature detection is therefore greatly diminished by missing values: a common problem in label-free DDA-MS, which relies on stochastic peptide sampling and MS/MS fragmentation of only the top ‘n’ peptides or other ions in a specified mass-to-charge ratio window (Jin et al., 2021; Webb-Robertson et al., 2015). Missing values are substantially reduced using DIA workflows such SWATH-MS (Dowell et al., 2021; Ludwig et al., 2018), for which both *CCprofiler* and PCprophet were primary developed. Further factors limiting protein complex identification in our dataset (compared with previous datasets benchmarked by *CCprofiler* and PCprophet) include separation of the cell lysate into a smaller number of SEC fractions, and the use of protein-level rather than peptide-level intensity data. Although we could have used the peptide-level intensities from MaxQuant for *CCprofiler* and PCprophet, we decided to avoid this approach in light of recent findings that peptide-level quantification results in significantly lower true positive rates than protein-level quantification for label-free DDA-MS data, especially with four replicates or fewer (Dowell et al., 2021).

The most informative experiment in assessing the impact of the various differences between our dataset and previously published datasets on protein complex identification has been provided by a SEC-MS study where the authors re-ran their samples using DDA instead of DIA (Heusel et al., 2019). Despite separating the lysate into three times as many SEC fractions as our study (81 *vs*. 24), and using peptide-level rather than protein-level intensities as the data input, *CCprofiler* identified 298 CORUM protein complexes from their DDA-SEC-MS dataset—only modestly more than our 245 protein complexes. Therefore, DIA appears to be the major contributor to the higher protein complex identifications in the more recent published SEC-MS studies. If DDA must be used, protein- or peptide-level labelling workflows (e.g., SILAC, ™T) could help to improve protein complex identification by reducing technical variability and/or missing values between conditions (Muntel et al., 2019; O’Connell et al., 2018; Stepath et al., 2020).

The limitation in protein complex detection also undoubtedly affected the ability to classify HSP90i-modulated complexes through PCprophet. Even among the 49 HSP90i-modulated complexes, we noticed that the majority of these had subunits whose abundances changed between the two conditions, without the remaining subunit profiles being drastically redistributed. If a protein complex was being disrupted as a whole, one would expect all the subunits of the complex to have at least one shifted peak (i.e., the co-elution peak for the disrupted complex). This interpretation was confirmed when we plotted the proteins differentially-regulated between Control and HSP90i conditions identified by PCprophet’s protein-centric workflow onto the protein complexes it identified through its complex-centric workflow (Fig S5). Very few complexes—aside from those with only two subunits—had the majority of their constituent subunits identified as differentially regulated. Therefore, we were unsure how many of our HSP90i-modulated complexes were truly being disassembled following HSP90 inhibition. This was one of the major rationales for us proceeding with the individual fraction-level differential analysis described in Fig 4. Indeed, the IDH3 complex SEC profiles demonstrated a more typical signature of disassembly, i.e., a significantly reduced peak, at the expected molecular weight of the functional complex, for all three constituent subunits following HSP90 inhibition (Fig 6A).

With more studies employing SEC-MS and other cofractionation-based MS approaches, we expect our dataset (e.g., Table S2) to continue being of use as newer analysis pipelines, software, and best-practices are developed (Schlossarek et al., 2021; Pang et al., 2020; Serwetnyk and Blagg, 2021). We also hope our study lays the foundation for other cutting-edge mass spectrometric approaches that increase resolution of treatment-induced changes to protein complexes or organellar proteomes, including SEC-SWATH-MS (Heusel et al., 2020), the various LOPIT techniques (Geladaki et al., 2019; Mulvey et al., 2017), and Dynamic Organellar Maps (Itzhak et al., 2016). A combination of these approaches could prove especially useful to test the epi-chaperome concept (i.e., that cancer cells are more dependent on highly inter-connected chaperone networks than non-cancer cells) with respect to HSP90 inhibition (Wang et al., 2019).

Another useful future comparison would be SEC-MS with other HSP90 inhibitors—especially those that bind HSP90 outside of its N-terminal ATPase pocket, such as C-terminal domain binders of the novobiocin family (Donnelly and Blagg, 2008) or covalent inhibitors targeting cysteine residues (Li et al., 2021; Zhu et al., 2021). C-terminal HSP90 binders represent an area of renewed interest (Armstrong et al., 2016; Bhatia et al., 2018; Goode et al., 2017; Park et al., 2020; Terracciano et al., 2018) due to their general propensity to inhibit chaperone activity without inducing the cytoprotective heat shock response—one of the major explanations proposed for the limited tumor-killing effect of N-terminal HSP90 inhibitors (Bickel and Gohlke, 2019). In a similar vein, inhibitors that target only a subset of the pan-HSP90 proteome by interfering with specific chaperone : co-chaperone interactions (e.g., HSP90 : CDC37 to target only protein kinase clients)(Serwetnyk and Blagg, 2021), or only certain HSP90 isoforms (Khandelwal et al., 2018; Park et al., 2020; Sanchez-Martin et al., 2020; Stothert et al., 2017), are also being explored for clinical use. The ability of SEC-MS to identify remodeling events downstream of HSP90 in a global and unbiased manner should make this an attractive approach for rational development and deployment of next-generation HSP90 family inhibitors to the patients and pathological indications most likely to benefit.

## Methods

### Cell culture

HT29 human colon adenocarcinoma, HCT116 human colon carcinoma, and BT474 human breast ductal carcinoma cells purchased from ATCC (LGC Promochem) were cultured in DMEM (Invitrogen) and supplemented with 10 % fetal calf serum (PAA Lab-oratories), 2 mm L-glutamine, 0.1 mm non-essential amino acids and 100 U of penicillin and streptomycin (all from Invitrogen) at 37 °C in a humidified incubator with 5 % CO_2_ and sub-cultured at 70 % confluency. Cells were confirmed as mycoplasma-free using the Venor Mycoplasma PCR Detection Kit (Minerva Biolabs), and were authenticated by short tandem repeat DNA profiling.

### Compound treatment and cell lysis for SEC

Five 15 cm dishes (50 % confluent) of HT29 cells were treated with 62.5 nm tanespimycin (Invivogen, CA, USA) (equivalent to 5x GI_50_ for the cell line), or mock-treated with equivalent volume of DMSO. The GI_50_ concentration for the cell line was determined by 96 h sulforhodamine B assay (Holford et al., 1998), and defined as the drug concentration that reduced the mean absorbance at 540 nm to 50 % of vehicle-treated controls. After 8 h, the cells were scraped on ice in 500 µL of ice-cold PBS containing cOmplete^™^ Protease Inhibitor Cocktail EDTA-free (Roche) and PhosStop (Roche). The collected cells were sonicated with a Diagenode Bioruptor (30 cycles: 30 s on, 30 s off) at 4 °C and then centrifuged at 17,000 *g* for 10 min at 4 °C. Samples were filtered through 0.45 µm Ultrafree-MC centrifugal filter units (Millipore) at 12,000 *g* for 10 min.

### SEC, enzymatic digestion, and peptide clean-up

Using a Dionex UltiMate 3000 HPLC system (Thermo Scientific), lysates were injected (200 µL per injection) onto a Superose^®^ 6 10/300GL column (GE Healthcare, UK) equilibriated with PBS (pH 7.2) with a flow rate of 0.2 mL min^−1^. 24 200 µL fractions were collected in a low protein-binding 96-deep-well plate (Eppendorf, Germany). Approximate protein concentrations were estimated using the EZQ^™^ Protein Quantitation Kit (Thermo Scientific). Tris-HCl (1 m, pH 8.0) was added to each fraction to a final concentration of 0.1 m Tris-HCl to adjust the pH to 8.0. After reduction and alkylation using DTT and iodoacetamide, respectively, proteins in each fraction were digested to peptides for 18 h at 37 °C using either trypsin alone or both LysC & trypsin diluted in 0.1 m Tris-HCl pH 8.0 at a final enzyme to protein ratio of 1:50 by weight. For peptide desalting, trifluoroacetic acid (TFA) was added to a 1 % (v/v) final concentration, and peptides were purified using a Sep-Pak tC18 96-well *µ*-elution plate (Waters). Peptides were eluted in 500 µL of 50 % (v/v) acetonitrile, 0.1 % TFA, and dried in a SpeedVac prior to resuspension in 5 % (v/v) formic acid. Peptide concentrations were determined using the CBQCA assay (Thermo Scientific) after 25-fold dilution of peptide samples in 0.1 m borate buffer, pH 9.3.

### LC-MS/MS and analysis of spectra

Using a Thermo Scientific Ultimate 3000 nanoHPLC system, 1 µg of peptides in 5 % (v/v) formic acid (approximately 10 µL) was injected onto an Acclaim^™^ PepMap^™^ C18 nano-trap column (Thermo Scientific). After washing with 2 % (v/v) acetonitrile in 0.1 % (v/v) formic acid, peptides were resolved on a 150 mm x 75 µm Acclaim^™^ PepMap^™^ C18 reverse-phase analytical column over a gradient from 2 % to 80 % acetonitrile over 100 min with a flow rate of 300 nL min^−1^. The peptides were ionized by nano-electrospray ionization at 1.2 kV using a fused silica emitter with an internal diameter of 5 µm (New Objective, Woburn, MA). Tandem mass spectrometry analysis was carried out on an LTQ Orbitrap Velos mass spectrometer (Thermo Scientific) using CID fragmentation. Data-dependent acquisition (DDA) involved acquiring MS/MS spectra on the 30 most abundant ions at any point during the gradient. The raw MS proteomics data have been deposited to the ProteomeXchange Consortium (http://proteomecentral.proteomexchange.org) via the PRIDE partner repository (Perez-Riverol et al., 2021) with the dataset identifier PXD033459.

Raw data were processed using MaxQuant software (http://www.coxdocs.org/doku.php?id=maxquant:start, version 1.5.1.3)(Cox and Mann, 2008) using the default settings and searched against the human UniProt database (June 7, 2011) with common contaminant entries. The settings used for MaxQuant analysis were: enzymes set as LysC/P and Trypsin/P, with maximum of 2 missed cleavages; fixed modification was Carbamidomethyl (Cys); variable modifications were Acetyl (Protein N-term), Carbamidomethyl (His, Lys), Carbamidomethyl (N-term), Deamidation (Gln, Asn), DiCarbamidomethyl (His, Lys), DiCarbamidomethyl (N-term), Gln to pyro-Gle, Oxidation (Met); Mass tolerance 20 ppm (FTMS) and 0.5 Da (ITMS); False Discovery Rate for both protein and peptide identification was 0.01. The ‘Re-quantify’ and ‘Match between runs’ features were both enabled. See Table S1 for the proteinGroups.txt output file from MaxQuant analysis.

Note that initial attempts to analyse the 384 raw files with MaxQuant indicated errors in reading five files from the EXP2 samples (trypsin-digested fractions 17, 22, & 24 in the Control condition, and LysC+trypsin-digested fractions 1 & 2 in the HSP90i condition). Therefore, for these fractions, only the sample digested with the other (MaxQuant-readable) enzyme schema was included in the MaxQuant input files.

### Initial SEC-MS data filtering and exploration

All data filtering, exploration, and statistical analyses were performed using a combination of Microsoft Excel and *R* (https://cran.r-project.org/, version 3.6.2) with Tidyverse, unless otherwise stated. Specific *R* packages are referenced in the following text. The proteinGroups.txt file from the MaxQuant analysis (Table S1) was used as the input data for all downstream statistical analysis reported here. Of the 7,401 entries in the protein-Groups.txt file, we filtered out 134 entries identified as potential contaminants, 91 as reverse matches, and 200 that were only identified by site. Additionally, 172 entries were removed as they were only identified by a single peptide. Of the remaining 6,804 proteinGroups entries, we consolidated into single entries the splice variants, duplicated Entrez protein IDs, and duplicated HUGO Gene Names—resulting in 6,427 unique protein entries for all subsequent analyses (Table S2). Gene Names that were not automatically assigned by MaxQuant were manually added from their Entrez protein IDs. To establish overlap and correlation between protein identifications across replicates, we used the *R* packages ‘UpSetR’ (Conway et al., 2017)(Fig 1C), ‘ggvenn’ (Fig S1A-B), and ‘heatmap.2’ (Fig S1C). For scaled intensities in heatmaps and linegraphs, LFQ fraction intensities were scaled (using ‘resca’ function from *R* package ‘metan’ (Olivoto and Lúcio, 2020))(Table S3). Scaling was performed per experiment (EXP1–EXP4) across all 48 fractions, such that the highest fraction intensity value for a protein in each EXP was set at 1, regardless of whether it was observed in the Control or HSP90i condition. For filtering based on number of replicates, we filtered for the proteins that had non-zero LFQ intensities in at least 1 of the 24 fractions (either Control or HSP90i condition—not necessarily both), in at least 3 of the 4 replicates, resulting in a list of 4,645 proteins (Table S4). Heatmaps in Figs 1D, 3D, and 4A were generated using the *R* packages ‘hclust’ for hierarchical clustering based on Euclidean distance with Ward-D2 linkage method and ‘heatmap.2’ for heatmap plotting. All boxplots, linegraphs, and Volcano plots were generated using the ‘ggplot2’ *R* package, unless otherwise stated.

### Limma-based differential expression analysis

All data shown at a total or summed level (i.e., without individual fraction values) have been processed in the following way. Starting with the list of 4,645 filtered proteins (Table S4), we added up all 24 LFQ intensities (F01:F24) for Control and HSP90i conditions separately. After log_2_-transforming and normalizing these data (variance-stabilizing normalization)(Fig S1D–E), we evaluated missing values, which were clearly biased towards proteins with lower LFQ intensities (Fig S1F). Additionally, there were a larger number of missing values in Experiment 1 (both Control and HSP90i)(Fig S1G). Based on these observations, we imputed the missing values using a manual left-censored Missing Not At Random (MNAR) method against a Gaussian distribution with a left-shift of 1.8 and a scale of 0.3. Using these parameters, we performed differential expression analysis using the *R* package DEP (https://bioconductor.org/packages/release/bioc/html/DEP.html) (Zhang et al., 2018) on the contrast between HSP90i and Control, with differentially-expressed proteins (DPs) set as those with an adjusted *p*-value of < 0.05 and an absolute log_2_FC threshold of 1 (i.e. Fold Change < -2 or > 2). The adjusted *p*-values depicted in summed boxplots for all figures are based on differential expression values calculated by this workflow. To calculate enriched Gene Ontology (GO) terms, we separately entered the significantly down-regulated (no enriched GO terms) and up-regulated (Fig 1H, Fig S1J) DPs into GOnet (Pomaznoy et al., 2018), using the full list of 4,645 filtered proteins as the background. Networks were visualized using CytoScape (version 3.8.2). For individual fraction-level differential analysis of the filtered 4,645 proteins (Fig 4, Table S9–S10), the same workflow was followed as for the summed protein intensities, except treating each of the 24 fractions as a separate Control *vs*. HSP90i DEP analysis, and without imputation of missing values. The DPs from each of the 24 fractions were combined, resulting in 366 DPs (Table S9). For the stringent fraction DPs (Table S10), only proteins that were DPs in two or more fractions were included. The network of stringent fraction DPs (Fig 4C) were generated using the STRING protein–protein interaction database (https://string-db.org/, version 11.5), with the following parameters: full STRING network type (both physical and functional interactions); edge thickness indicating strength of evidence for interaction; minimum required interaction score = 0.4 (medium confidence); Markov Clustering (MCL) with inflation parameter = 2. GO term enrichment analysis was also performed on STRING, using the full list of 4,645 filtered proteins as background. The canSAR curated interactome-based network (Fig S6) was generated using the canSAR Protein Annotation Tool (https://cansarblack.icr.ac.uk/cpat). The canSAR interactome contains > 1 million binary interactions for > 19,000 human proteins. Interaction types are classified to reflect the method of experimental determination. A confidence level in the existence of a direct binary interaction is assigned in canSAR based on the type of the experiment and the number of independent publications reporting the interaction. All experimental determination types were included with a confidence level of >= 0.1.

### *CCprofiler* and PCprophet protein complex detection

To identify the number of CORUM-annotated protein complexes preserved in our dataset, we used either *CCprofiler* (Heusel et al., 2019) or PCprophet (Fossati et al., 2021), with the data frame of 6,427 proteins used as a starting point (Fig S2). For *CCprofiler*, missing values in the computed list of protein traces were imputed by fitting a spline interpolation, and normalized by cyclic loess (Bludau et al., 2021). The proportion of intensities in assembled and monomer range (Fig 3A) were estimated using the *CCprofiler* ‘summarizeMassDistribution’ function. For feature detection purposes, protein traces were aggregated across conditions and replicates by summing up intensity values in each fraction. All subsequent complex-centric feature detection and co-elution (Fig 3B, Fig S3A–B) was performed against the default CORUM complexes reference data frame ‘corumComplexHypothesesRedundant’ in *CCprofiler*. Decoy complex queries were generated from this reference data frame (min_distance = 2), and protein complex features were detected from our normalized protein traces with the following parameters: corr_cutoff = 0.9, window_size = 5, rt_height = 1, smoothing_length = 5, collapse_method = “apex_network”, perturb_cutoff = 5 %. The resultant complex features were filtered according to their apparent molecular weight (min_monomer_distance_factor = 1.2). The complex co-elution peak groups with the largest number of co-eluting protein subunits (‘getBestFeatures’ function) were then selected for statistical scoring at a 5 % FDR. For PCprophet, data from Table S2 was separated into 8 separate txt files (Ctrl_1–Ctrl_4, HSP90i_1–HSP90i_4), together with a sample ID key and calibration table for molecular weight estimation, as outlined in PCprophet instructions (https://github.com/anfoss/PCprophet/blob/master/PCprophet_instructions.md). PCprophet was then run (using Python v3.7.3) on this dataset with default parameters against the CORUM database (using the ‘coreComplexes.txt’ file included in PCprophet), except with calibration by molecular weight (-cal) turned on, mapping of gene names to molecular weight (-mw_uniprot) included as a file from UniProt, and molecular weight-based complex collapsing (-co CAL flag). From the PCprophet output, the ‘DifferentialProteinReport.txt’ file was used to identify differential proteins with a ‘Probability_differential_abundance’ > 0.5 (Fig 3E, Table S7), and a combination of the ‘ComplexReport.txt’ and ‘DifferentialComplexReport.txt’ files to identify positive complexes (‘Is Complex’ = Positive) and differential complexes (‘Is Complex’ = Positive AND ‘Probability_differential_abundance’ > 0.5)(Fig 3E, Table S8). The SEC profiles of all subunits identified by PCprophet in different complexes were plotted as linegraphs (Fig 3F–G, Fig S4). The protein–protein interaction networks (Fig S5) were generated by importing ‘PPIReport.txt’ PCprophet output into CytoScape (colour = Control or HSP90i; edge width = count(‘Replicate’) grouped by Control or HSP90i), and differential proteins from Table S8 were mapped onto the network nodes.

### SEC-Immunoblotting

The cell culture, compound treatment, cell lysis, and SEC protocols used for SEC-MS were followed as closely as possible for validation by SEC-IB. Five 15 cm dishes (80 % confluent) of HT29 cells were treated with 62.5 nm tanespimycin (equivalent to 5 x GI_50_ for the cell line), or mock-treated with equivalent volume of DMSO. After 8 h, the cells were scraped on ice in 500 µL of ice-cold PBS containing cOmplete^™^ Protease Inhibitor Cocktail EDTA-free (Roche) and Phosphatase Inhibitor Cocktails 1 & 2 (Sigma). The collected cells were sonicated with a Branson-Tip Sonicator (high power, 3 cycles: 30 s on, 30 s off) at 4 °C and then centrifuged at 17,000 x *g* for 10 min at 4 °C. Samples were filtered through 0.45 µm Ultrafree-MC centrifugal filter units (Millipore) at 12,000 x *g* for 10 min. BCA protein assays (Pierce) were performed on the filtrates for protein quantification.

Using an ÄKTApurifier UPC 10 FPLC system (GE Healthcare), lysates in PBS with protease and phosphatase inhibitor cocktails were injected (500 µL per injection, corresponding to 1–3 mg total protein) onto a Superose^®^ 6 10/300GL column (GE Life Sciences) equilibriated with PBS (pH 7.2) with a flow rate of 0.2 mL min^−1^. After 10 mL of void volume, 500 µL fractions were collected using a low protein-binding 96-deep-well plate (Eppendorf). Fractions were aliquoted and stored at -80 °C before adding 3X Blue Loading Buffer (Cell Signaling Technologies) to 25 µL of each fraction, and running on NuPAGE 4–12 % Bis-Tris gels (Invitrogen). Following gel transfer onto nitrocellulose (Invitrogen), membranes were blocked in Tris-Buffered Saline (50 mm Tris-HCl, pH 7.5, 150 mm NaCl) with 1 % Tween-20 (TBS-T) supplemented with 5 % BSA (for HOP immunoblotting) or milk powder (for all other antibodies) for 1 h before incubating with the appropriate concentration of primary antibody diluted in TBS-T with BSA or milk powder overnight. Antibodies for HSP90α/β (sc-7947, 1:500), CDC37 (sc-5617, 1:500), and AHA1 (sc-50527, 1:500) were from Santa Cruz Biotechnology; p23 (ab92503, 1:10,000) and Anillin (ab99352, 1:2,000) from Abcam; BAG3 (10599-1-AP, 1:1,000) from Proteintech; Hop (#4464, 1:2,000) from Cell Signaling Technologies; and HSP70/HSP72 (ADI-SPA-810-D, 1:2,500) from Enzo Life Sciences. Membranes were then washed with TBS-T (3x, 5 min each) and incubated with horse radish peroxidase (HRP)-conjugated secondary antibodies (GE Healthcare). Following another wash step in TBS-T, the HRP signal was detected by incubation with Pierce^™^ ECL Western Blotting Substrate (Thermo Scientific) and exposure to Hyperfilm ECL (GE Healthcare). Immunoblots shown are representative of three independent experiments.

For the HSP90 co-immunoprecipitation experiment shown in Fig 5B, 1 mg of lysate from HT29 cells treated with tanespimycin or mock vehicle control as described above was diluted in modified RIPA buffer (50 mm Tris-HCl pH 7.5, 150 mm NaCl, 1 % IGEPAL CA-630, 0.5 % sodium deoxycholate, 0.02 % SDS, cOmplete^™^ Protease Inhibitor Cocktail EDTA-free (Roche)) to 190 µL final volume and incubated with 10 µL anti-HSP90 conjugated magnetic beads (clone SJ-90; LSBio catalog no. LS-C171164) for 1 h at 4 °C under rotary agitation. Two negative controls were incubated in parallel: one with a 50:50 mix of the tanespimycin and mock-treated HT29 lysates (1 mg total protein) in the presence of control magnetic beads without any conjugated antibody (LifeSensors, catalog no. UM400M) (‘–IgG’), and a second control with the anti-HSP90 magnetic beads incubated with modified RIPA buffer only (‘+IgG’). Following five washes with modified RIPA buffer to remove unbound proteins, co-immunoprecipitated proteins were eluted from the magnetic beads by incubating with 10 µL 4X LDS Sample Buffer (Abcam) at 70 °C for 10 min, followed by transfer of the eluate into a separate tube and a further 10 min incubation at 70 °C in the presence of DTT (50 mm final concentration) to reduce the eluted proteins fully. Eluted proteins were loaded onto separate gels for each individual immunoblot. SDS-PAGE and immunoblotting were performed as described above, except with rabbit anti-HSP90 clone C45G5 (Cell Signaling Technologies #4877, 1:2,000) for the HSP90 immunoblot, and a 1:500 dilution (instead of 1:2,000) for the Anillin immunoblot.

See Fig S9 for uncropped images of all immunoblots displayed in this manuscript.

### Anillin knockdown experiments

siPools (siTools Biotech GmbH) comprising 30 siRNAs targeting Anillin (ANLN), Polo-like kinase 1 (PLK1), or no recognised mammalian gene (Non-targeting control, NTC), or sterile water (untargeted control, UTC), were complexed for 15 min with DharmaFECT 4 transfection lipid (Horizon Discovery) in 75 µL/well OptiMEM (Invitrogen) at room temperature. Following the 15 min incubation, 2,000 HT29 cells in 75 µL DMEM (without Penicillin/Streptomycin) were added to each well, giving a final concentration of 25 nm siRNA and 0.8 % lipid. Cells were incubated at 37 °C, 5 % CO_2_ in an Incucyte^®^ Zoom (Sartorius AG) and scanned every 8 h for 2 days. After 2 days, 75 µL tanespimycin (final concentrations from 0.137–300 nm) or DMSO (0.1 % final concentration) was added to each well. Cells continued to be scanned every 8 h for a further 4 days before being assessed for the number of viable cells using Cell Titer Blue^®^ (Promega) according to the manufacturers instructions. Cellular viability and confluency at each concentration of tanespimycin were calculated as a percentage of the DMSO control for the re-spective siRNA treatment with the dose response curves plotted as non-linear functions with variable slope in GraphPad Prism v. 9.0.0 (GraphPad Software). The effect of siRNA knockdown on cells was calculated as a percentage of the NTC. All experiments were performed in biological triplicate.

### Isocitrate dehydrogenase activity assays

IDH activity was measured using the Isocitrate Dehydrogenase Activity Assay Kit (Sigma catalog no. MAK062) according to the manufacturer’s protocol, with NAD^+^ or NADP^+^ used as the co-factor for IDH3 or IDH1/2, respectively. Briefly, 1 million HT29, HCT116, or BT474 cells were scraped on ice in 200 µL ice-cold IDH Assay Buffer (Sigma, Catalog Number MAK062A), centrifuged at 13,000 x *g* for 10 min at 4 °C. 50 µL of this cell lysate was added to an equal volume of the manufacturer’s Master Reaction Mix, with NAD^+^ or NADP^+^ for quantification of IDH3 or IDH1 & IDH2 activity, respectively. Absorbance readings at 450 nm for the samples compared with the NADH standard curve (always run in parallel with the samples) were used to estimate IDH activity.

### Manuscript preparation

This manuscript was prepared in Overleaf (http://www.overleaf.com), using the RoyleLab bioR*χ*iv template.

## ACKNOWLEDGEMENTS

We thank all colleagues for help with various aspects of the study, including Costas Mitsopoulos and Felix Krueger for computational set-up of the data analysis, Craig McAndrew for help with the FPLC for SEC-IB validation experiments, and Arne Smits for providing a comprehensive vignette on Bioconductor for DEP package use (https://www.bioconductor.org/packages/devel/bioc/vignettes/DEP/inst/doc/DEP.html). PW acknowledges grant funding from Cancer Research UK (Program Grants C309/A31322 and C309/A11566; Strategic Award C35696/A23187). PW is also funded by Wellcome Trust, Chordoma Foundation, and Mark Foundation, and is a Cancer Research UK Life Fellow. BAL is currently receiving funding from the Cancer Prevention and Research Institute of Texas, Cancer Research UK (Strategic Award C35696/A23187), and The Wellcome Trust. The mass spectrometry-based proteomics was funded by a supplement to Cancer Research UK Grant C309/A10705 to RSS and PW. SB was funded by Cancer Research UK Program Grants C309/A31322 and C309/A11566. RSS and HEJ are funded by Institute Strategic Programme Grant BB/P013384/1 from the BBSRC.

## AUTHOR CONTRIBUTIONS

**Conceptualization:** RSS, AIL, PAC, PW. **Data curation:** RSS, SB, ML, BO, CM, IB, LB, SA, BAL. **Formal analysis:** RSS, SB, ML, BO, CM, IB, BAL. **Funding acquisition:** RSS, PW. **Investigation:** RSS, SB, ML, BO, CM, IB, AH, HEJ. **Methodology:** RSS, SB, ML, BO, IB, SA, BAL, AIL, PAC, PW. **Project Administration:** RSS, AIL, PAC, PW.

**Resources:** RSS, SA, BAL, AIL, PW. **Software:** n/a **Supervision:** RSS, AIL, PAC, PW. **Validation:** RSS, SB, CM, AH, HEJ. **Visualization:** RSS, SB, BO, IB, LB, BAL. **Writing– original draft:** RSS. **Writing–review & editing:** RSS, SB, ML, BO, CM, IB, LB, SA, AH, HEJ, BAL, AIL, PAC, PW.

## Supplementary Information

### LIST OF SUPPLEMENTARY FIGURES & TABLES

**Figure S1.**
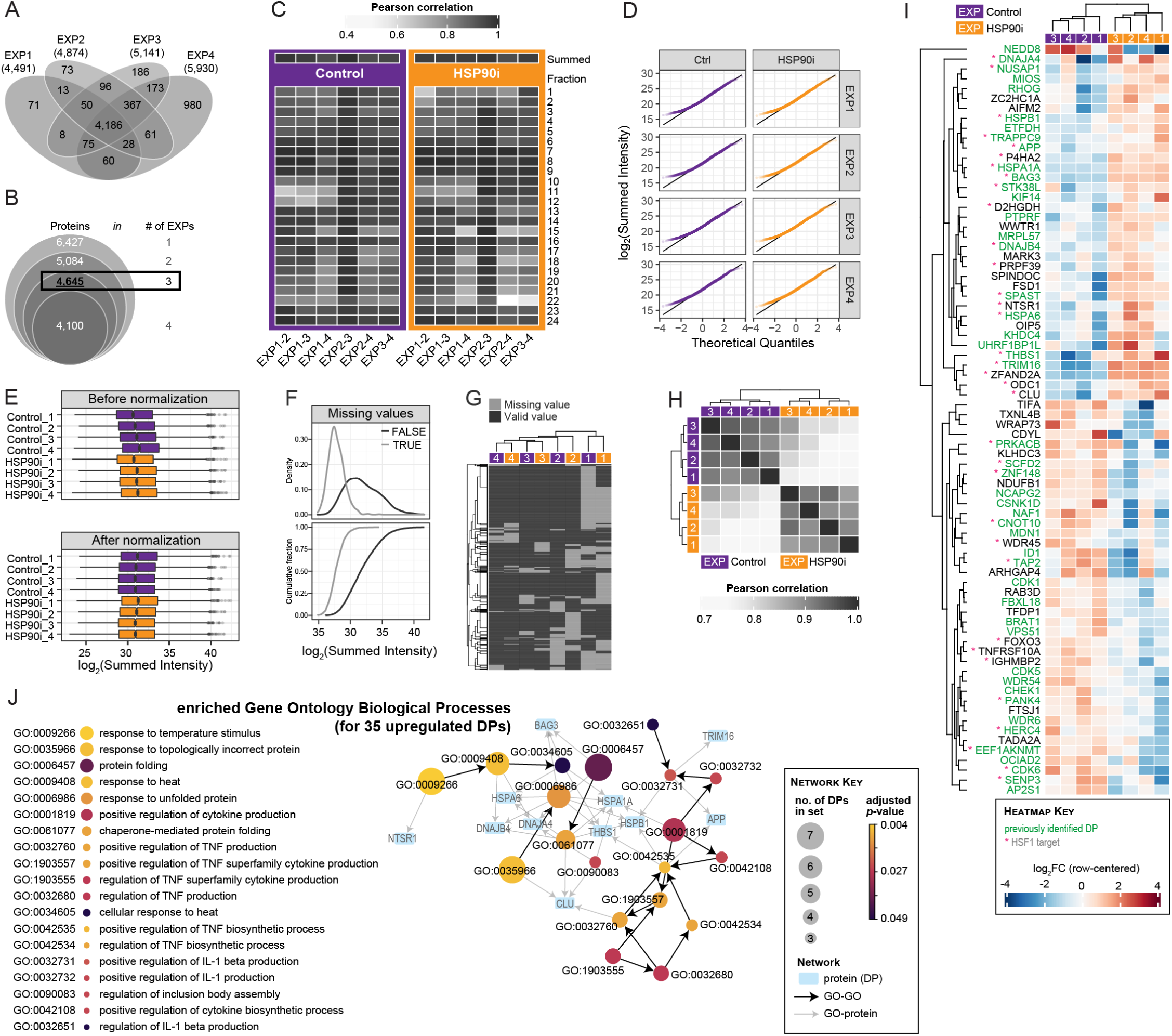
Reproducibility between replicates (EXP1–EXP4) and differential protein analysis of summed intensities in SEC-MS dataset. (**A**) Venn diagram showing overlap between the proteins identified in each biological replicate (EXP1–EXP4). (**B**) Venn diagram to show total number of proteins identified in 1, 2, 3, or 4 biological replicates. For (A) and (B), proteins identified in either Control or HSP90i conditions were combined. (**C**) Heatmap showing Pearson correlation coefficients between the four biological replicates for each of the 24 SEC fractions, individually and summed, under both experimental conditions. (**D**) Quantile-quantile plots of log_2_-transformed summed LFQ intensities from filtered dataset (4,645 proteins) indicate a similar heavy-tailed distribution in all experiments. This type of distribution is typical in mass-spectrometry-based proteomics data. (**E**) Box/Tukey plots showing spread of summed intensities for each experiments before and after background correction and variance-stabilizing transformation. (**F**) Linegraphs showing densities (top) and cumulative fractions (bottom) of log_2_-transformed summed intensities for proteins with (TRUE) and without (FALSE) missing values. Data indicate that proteins with missing values tend to have lower summed intensities. (**G**) Heatmap of entries from 4,645 filtered proteins list with at least one missing value across the eight samples. (**H**) Heatmap summarizing Pearson correlation coefficients for summed intensities across all experiments. (**I**) Heatmap showing row-centered intensities for all 76 significant differential-expressed proteins based on summed intensity values. Clusters are drawn from Euclidean distance. Heatmap and plots in (D)–(G) were generated from built-in functions in *R* package ‘DEP’. (**J**) Gene Ontology Biological Processes (GOBPs) significantly enriched among the 35 upregulated differential proteins (DPs) identified in Fig 1F using GOnet, with the 4,645 filtered proteins used as the background for the enrichment analysis. DPs linked to the enriched GO terms are shown.

**Figure S2.**
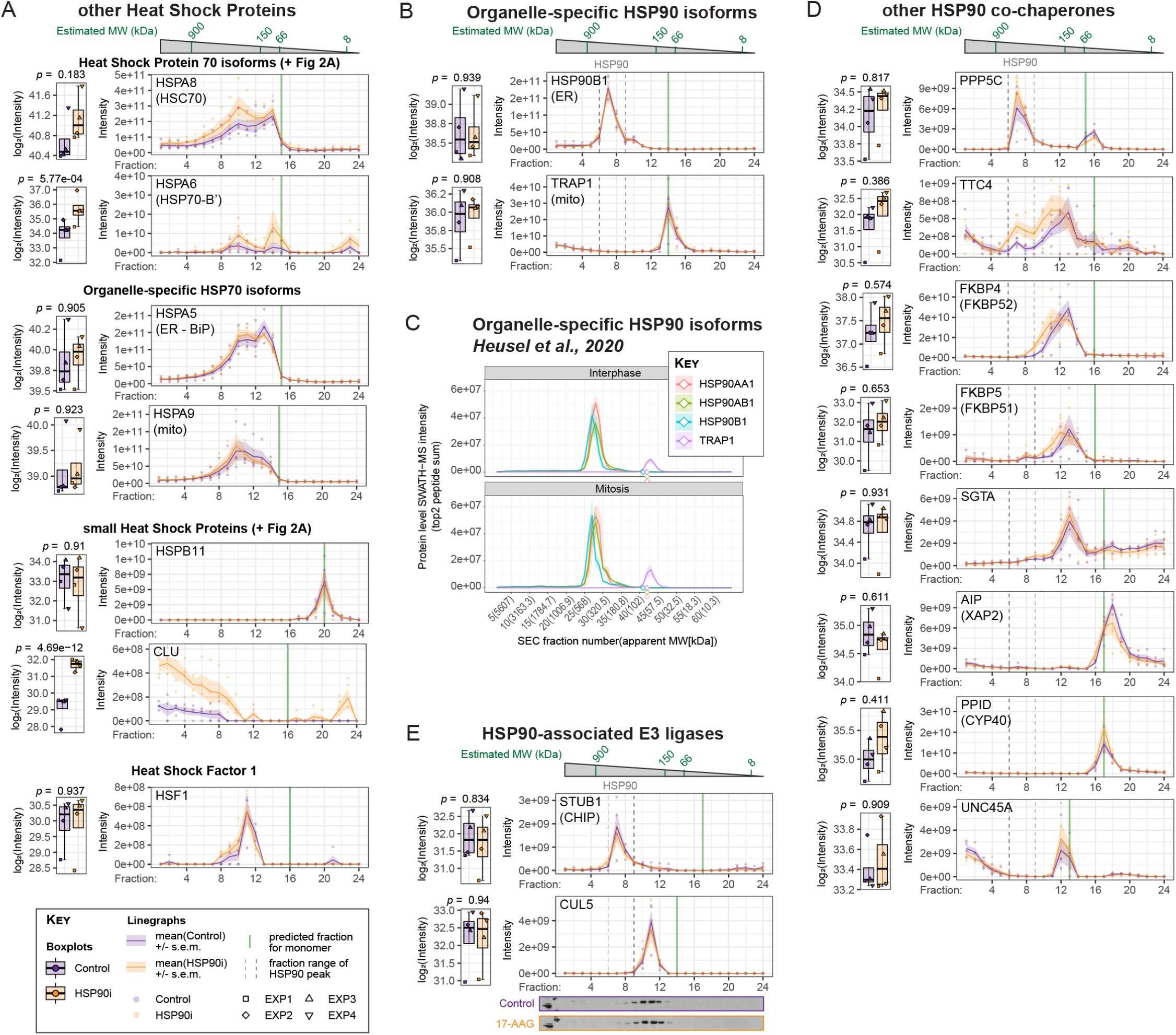
SEC-MS profiles of components of the Heat Shock Protein machinery. (**A**) SEC-MS profiles of other heat shock-induced proteins and their transcriptional activator HSF1. (**B**) SEC-MS profiles of organelle-specific HSP90 isoforms HSP90B1/GRP94 (endoplasmic reticulum) and TRAP1 (mitochondria). (**C**) SEC-SWATH-MS profiles from Heusel et al., 2020, for HSP90 isoforms HSP90AA1/HSP90*α*, HSP90AB1/HSP90*β*, HSP90B1/GRP94 (endoplasmic reticulum) and TRAP1 (mitochondria), from interphase and mitotic HeLa-CCL2 cells. Profiles were generated using SECexplorer-cc (https://sec-explorer.shinyapps.io/hela_cellcycle/). Solid lines represent mean values for the top two peptides across three replicated for each protein. Shaded areas represent the standard error of the mean. Diamonds indicate estimated fraction for the monomer peak of each protein, based on the UniProt-annotated molecular weight. (**D**) SEC-MS profiles of other tetratricopeptide (TPR)- domain co-chaperones of HSP90. Only PPP5C/PP5 had a major elution peak in the same fractions as HSP90. (**E**) SEC-MS profiles of two E3 ubiquitin ligases known to be recruited to HSP90 following HSP90 inhibition. Only STUB1/CHIP co-eluted with HSP90. Neither E3 ubiquitin ligase displayed an altered profile following HSP90 inhibition.

**Figure S3.**
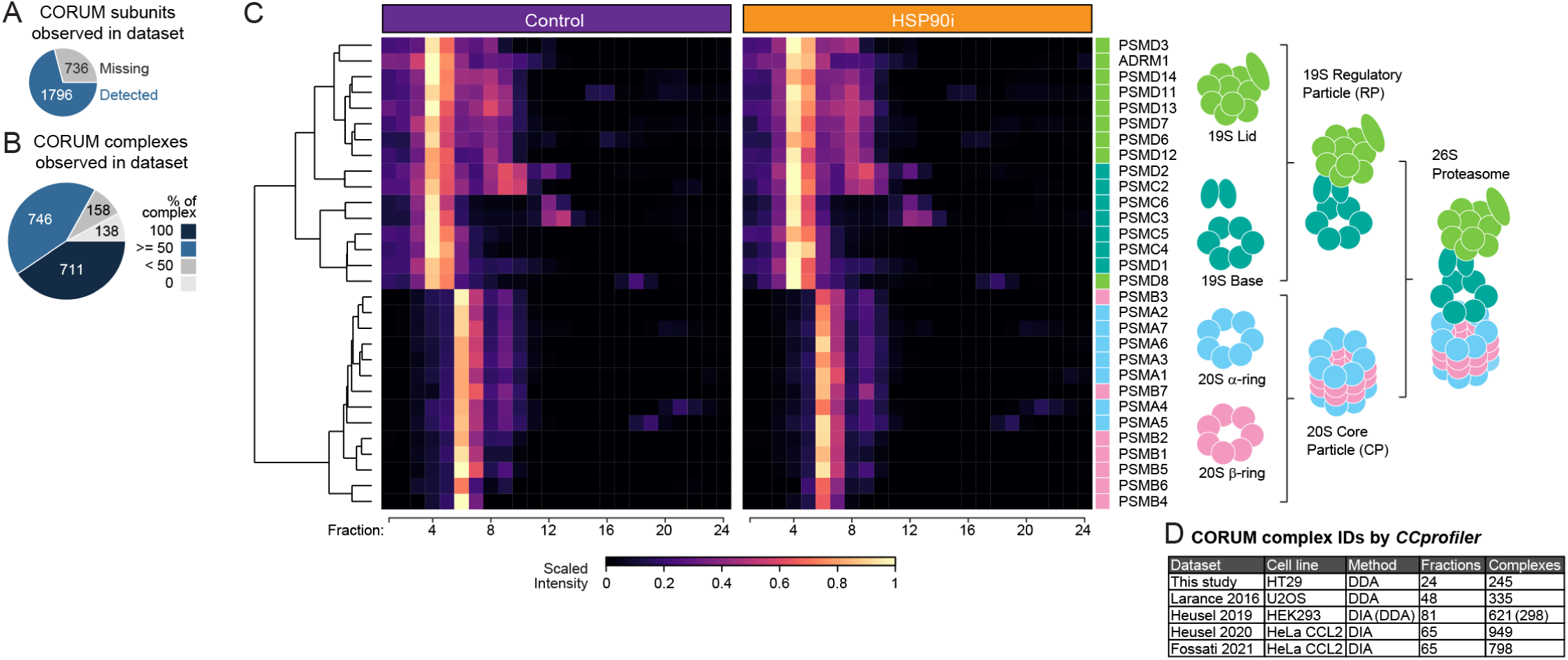
Tight clustering of proteins within, but not between, regulatory particle (RP, 19S) and core particle (CP, 20S) of the 26S proteasome. (**A–B**) Pie charts illustrating the number of proteins (A) and protein complexes (B) annotated in the CORUM protein complex database that were covered in our dataset of 6,427 proteins. (**C**) Heatmap of scaled intensities for subunits of the 19S regulatory particle and 20S core particle complexes that make up the 26S proteasome. The dendogram cut-offs based on Euclidean distance matrix with the Ward-D2 linkage method are illustrated to the left of the heat map. (**D**) Benchmarking the number of CORUM-annotated protein complexes identified by *CCprofiler* from our dataset compared with previous published SEC-MS studies. The cell line, MS acquisition method (data-dependent acquisition, DDA, or data-independent acquisition, DIA), number of SEC fractions collected, and number of protein complexes identified by *CCprofiler* (using a cut-off of >= 50 % of subunits co-eluting) are indicated. The number of complexes identified are according to those reported in each study, except for Larance et al., which pre-dates *CCprofiler*, and was re-analyzed using *CCprofiler* in Heusel et al. 2019.

**Figure S4.**
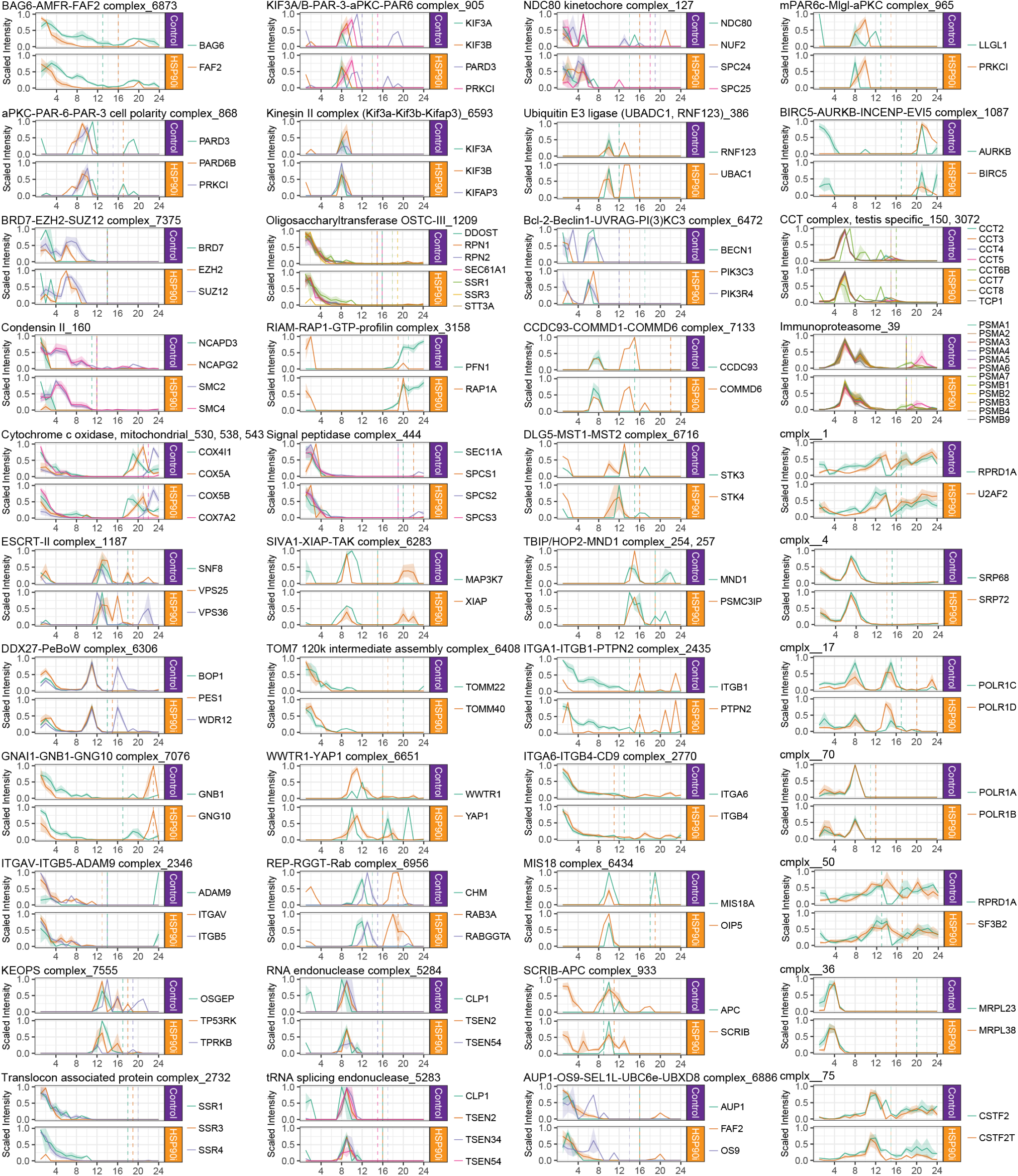
SEC profiles of all HSP90i-modulated protein complexes identified by PCprophet. Scaled median intensities for all subunits from the protein complex identified as co-eluting by PCprophet were plotted. CORUM ID numbers are indicated in the title. For complexes with multiple CORUM IDs, only one graph was plotted (e.g., ‘Cytochrome c oxidase, mitochondrial’ CORUM IDs 530, 538, and 542; ‘CCT complex, testis specific’ CORUM IDs 150 and 3072). Novel protein complexes not annotated in CORUM are indicated with the prefix “cmplx”. Dashed vertical lines on linegraphs indicate the fraction in which the monomer would be detected, based on the UniProt-annotated molecular weight.

**Figure S5.**
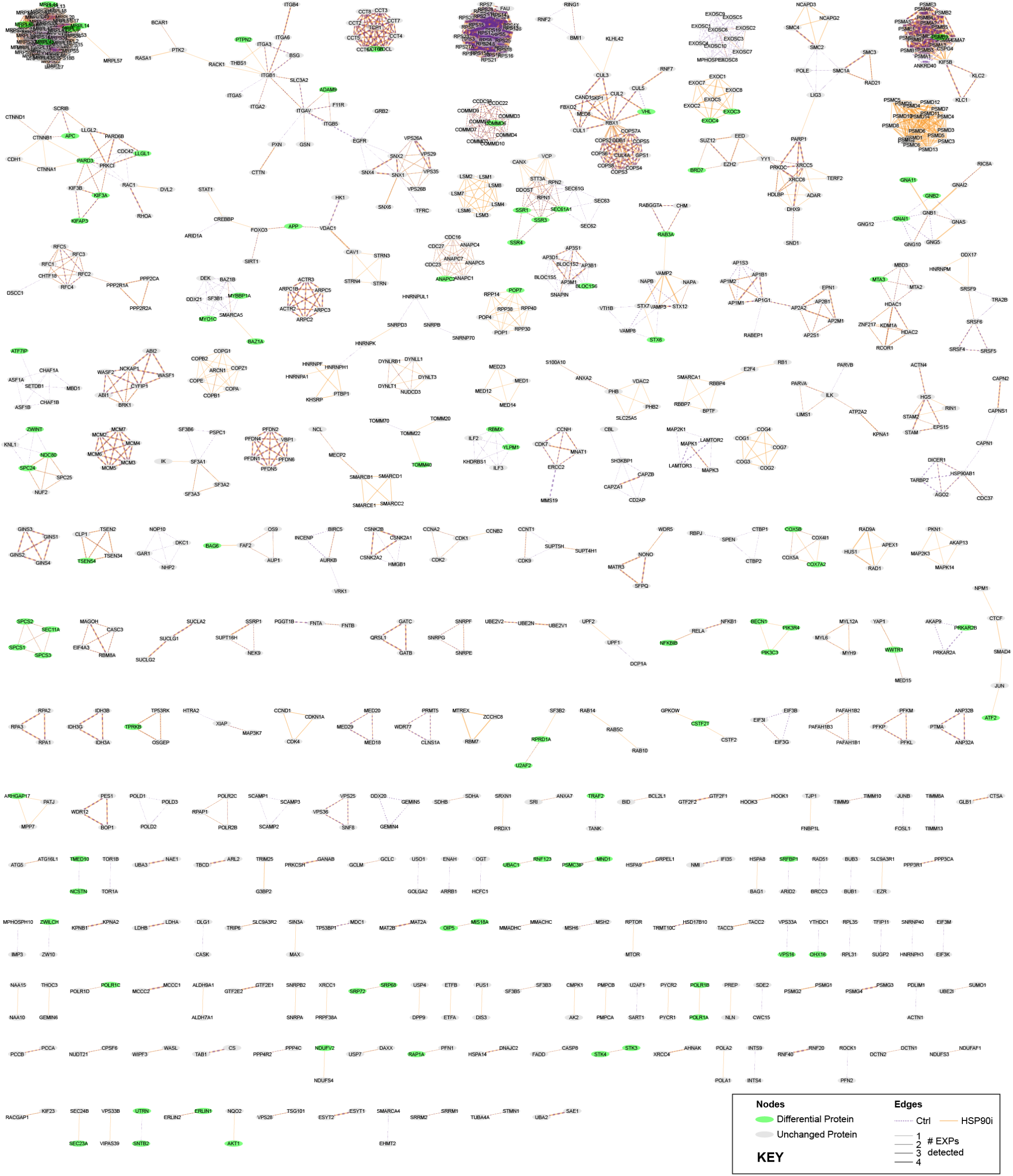
Protein–protein interactions networks identified in complete dataset. Protein–protein interaction networks generated by PCprophet based on our complete dataset of 6,427 proteins. Networks were generated using Cytoscape based on ‘PPIreport.txt’ PCprophet output. Green nodes represent differential proteins identified using PCprophet’s protein-centric analysis. Edge width represents the number of experiments in which the interaction was confidently detected by PCprophet, with the edge colour representing Ctrl or HSP90i detections.

**Figure S6.**
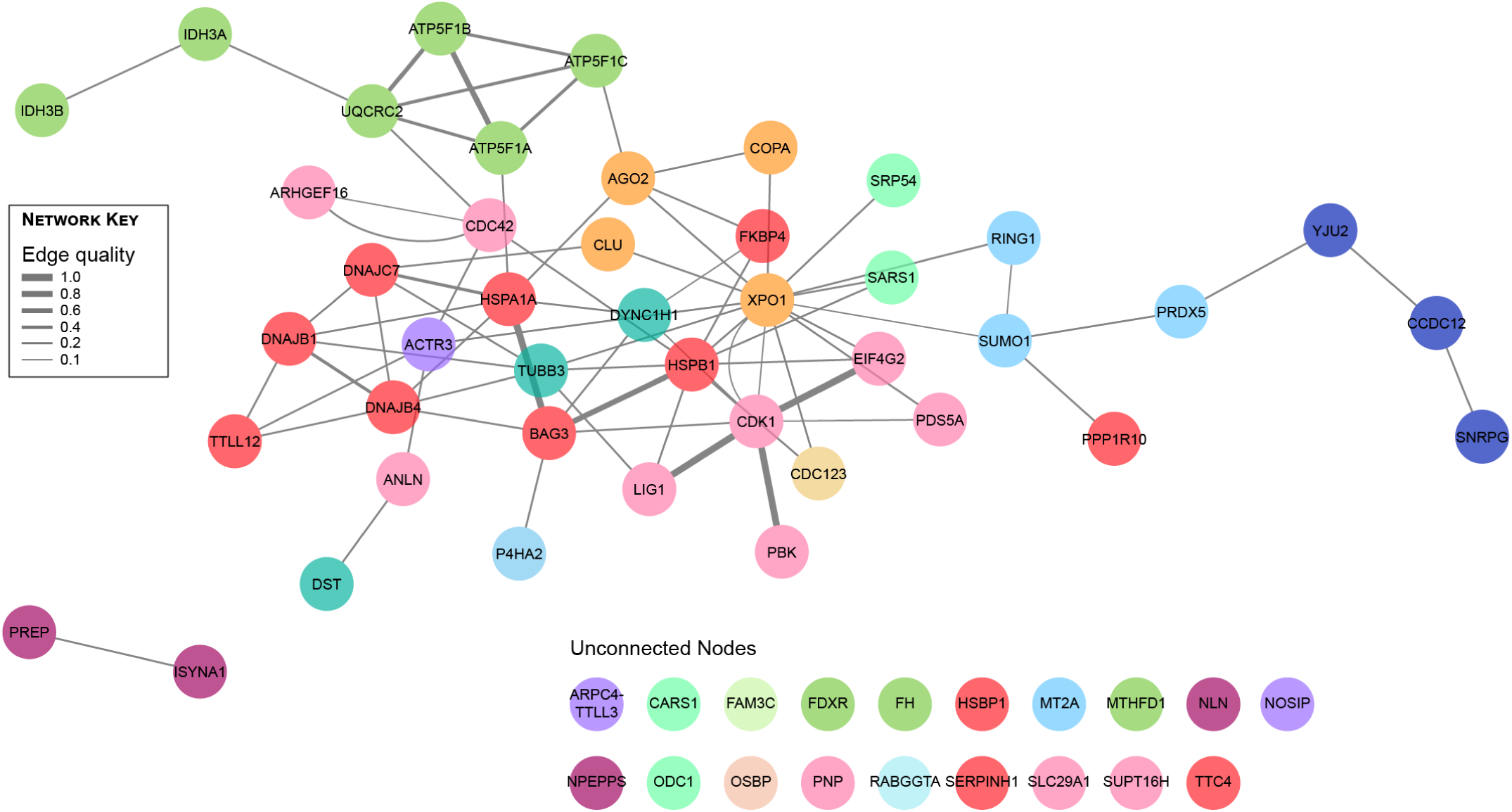
Stringent DP protein–protein interaction network generated using canSAR curated interactome database. The list of 62 stringent fraction DPs from Fig 4 were entered into the canSAR Protein Annotation Tool (https://cansarblack.icr.ac.uk/cpat), with edges representing interactions from canSAR’s curated interactome. Edge width represents confidence of evidence (‘edge quality’) for interaction between the two nodes. Node colours represent clusters from Fig 4.

**Figure S7.**
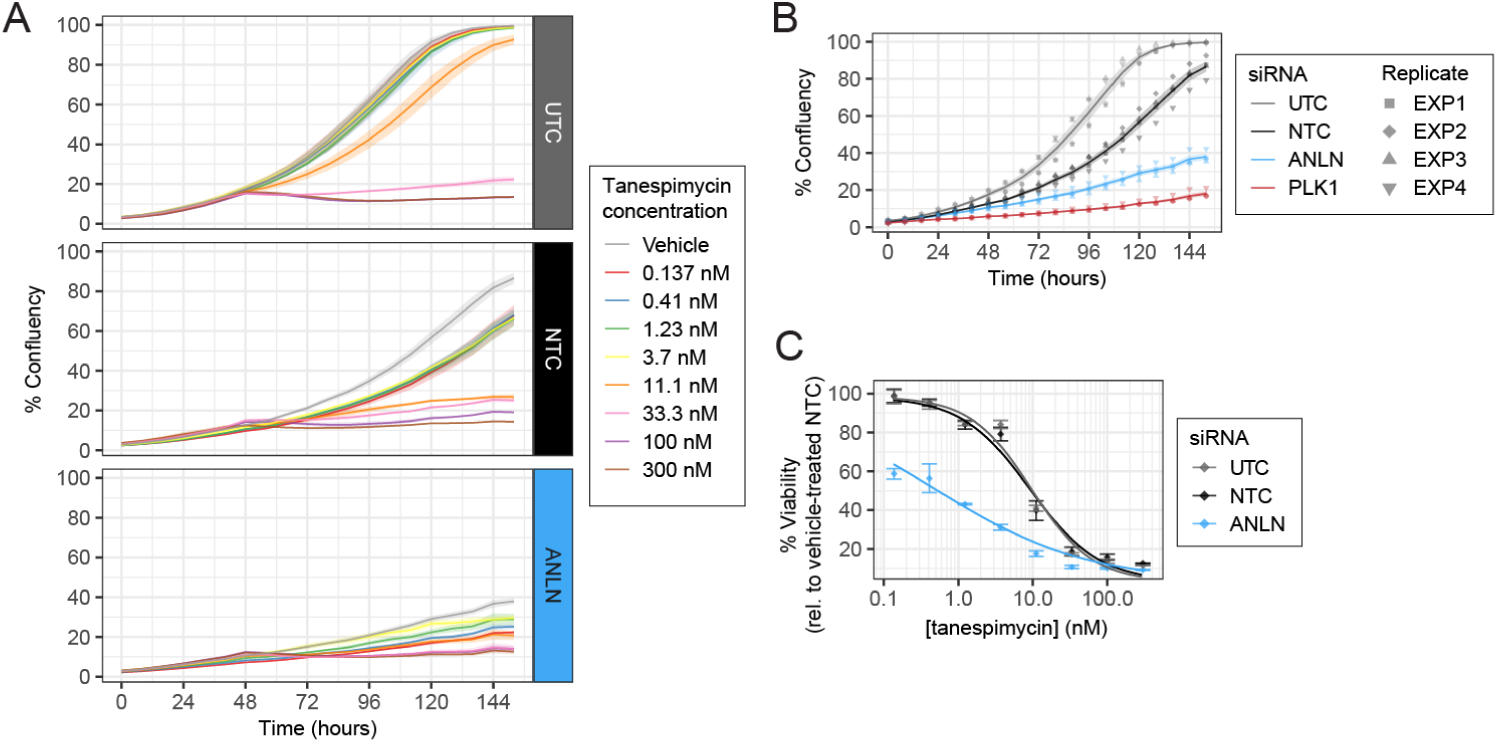
Knockdown of Anillin by siRNAs reduces cell confluency and viable cell number. (**A, B**) Knockdown of Anillin reduces HT29 cell confluency compared with non-targeting control (NTC) and untargeted control (UTC). HT29 cells were treated for 48 h with siTOOLs pool of 30 siRNAs (25 nM total concentration) targeted against Anillin (ANLN), non-targeting control NTC, untreated control UTC, or death-inducing control PLK1, followed by 96 h with a range of tanespimycin concentrations (except for PLK1), or with mock treated with vehicle control (0.1 % DMSO), while still in the presence of the original siRNAs. Confluency was monitored every 8 h throughout the time-course by Incucyte^®^. Lines and shaded regions represent mean +/- standard error from four biological replicates (EXP1–EXP4) for each condition. (**B**) Confluency measurements for vehicle control-treated cells from each siRNA condition in **A**. (**C**) Knockdown of Anillin reduces number of viable HT29 cells compared with non-targeting control (NTC) and untargeted control (UTC). Percentage viability of HT29 cells from part **A** at the end of the siRNA and tanespimycin treatments (144 h total) by CellTiter-Blue^®^ assay relative to vehicle-treated (0.1 % DMSO) NTC control was measured with an Incucyte^®^. Points and error bars represent mean +/- standard error for each condition. Dose-response curve fitting was performed using the ‘Log[Inhibitor] *vs*. normalized response – Variable slope’ non-linear regression model in Graphpad Prism.

**Figure S8.**
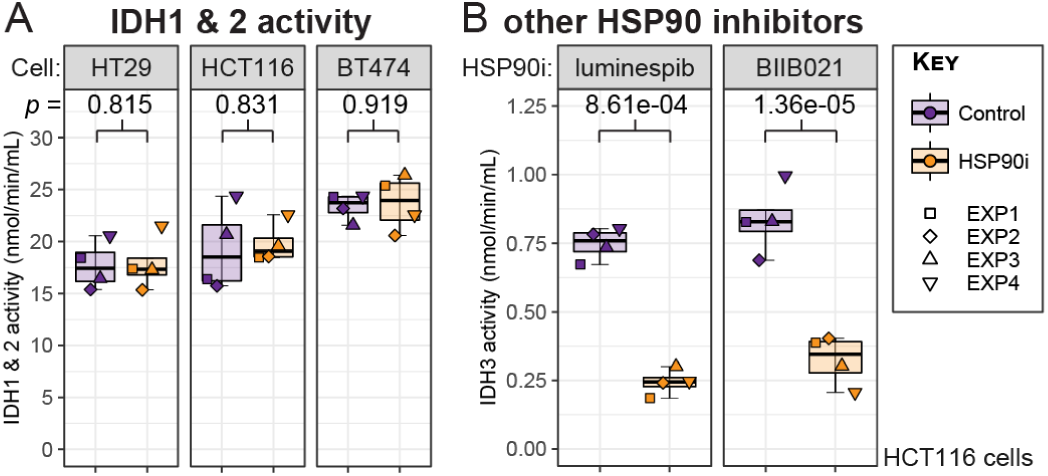
Activity of the mitochondrial Isocitrate Dehydrogenase 3 (IDH3) complex, but not IDH1 or IDH2, is reduced by treatment with multiple HSP90 inhibitors. (**A**) Combined activities of IDH1 & IDH2 are not significantly reduced upon HSP90 inhibition in HT29 colon adenocarcinoma, HCT116 colon carcinoma, and BT474 breast ductal carcinoma cell lines. Activity was measured using the IDH Activity Assay Kit (Sigma) according to manufacturer’s instructions, using NADP^+^ as the co-factor—which is used by both IDH1 and IDH2. See Fig 6B for corresponding assay with NAD^+^ as the co-factor, for estimating IDH3 activity. (**B**) IDH3 activity is significantly reduced upon treatment with the chemotypically distinct HSP90 inhibitors luminespib (AUY922) and BIIB021, in HCT116 colon carcinoma cells. Activity was measured using the IDH Activity Assay Kit (Sigma) according to manufacturer’s instructions, using NAD^+^ as the co-factor.

**Figure S9.**
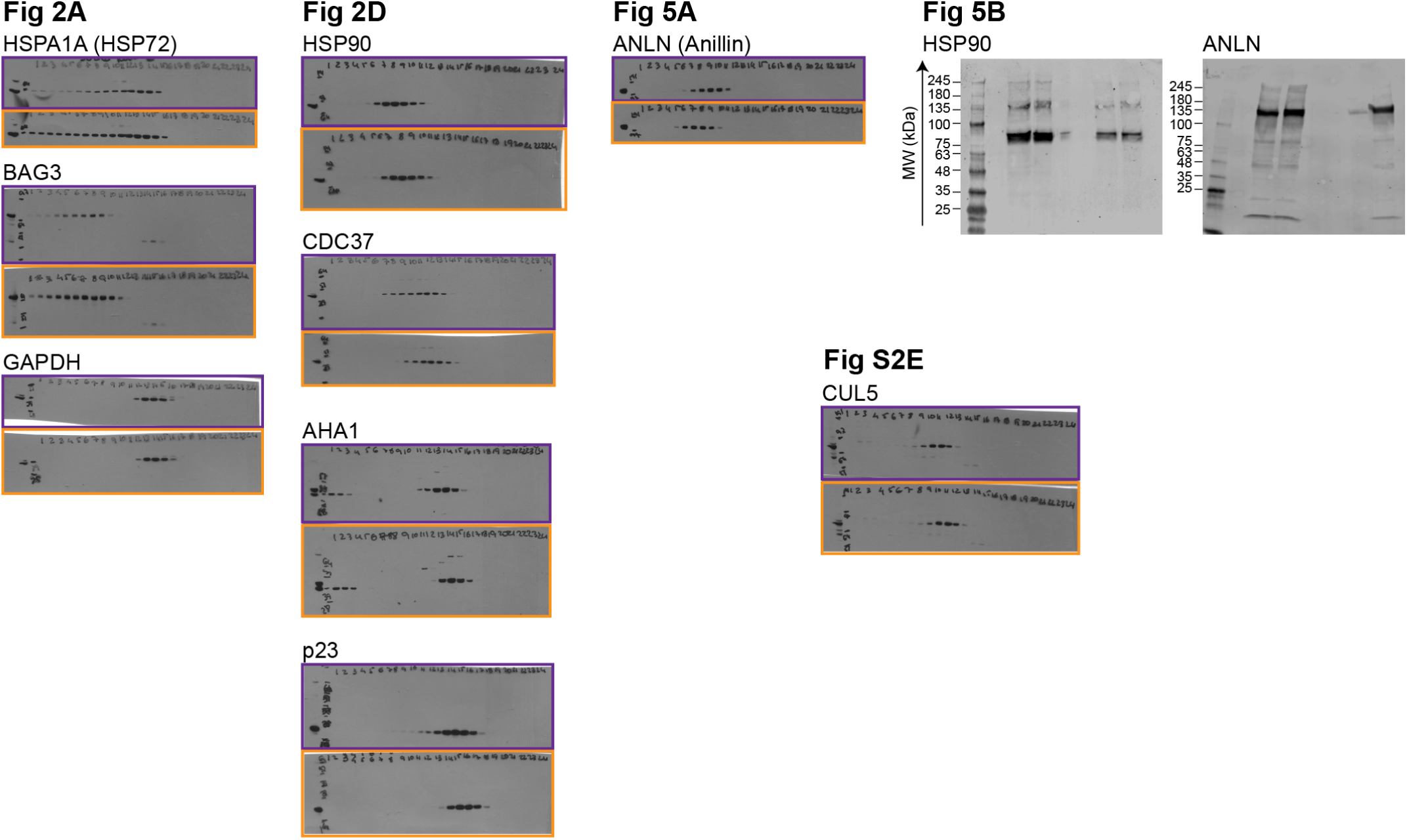
Uncropped images of immunoblots displayed in this manuscript. See ‘SEC-Immunoblotting’ section in Methods for antibodies and dilutions used.

### SUPPLEMENTARY TABLE LEGENDS

**Table S1. Original proteinGroups.txt file from MaxQuant**. Data-frame of 7401 rows and 1595 columns. See MaxQuant documentation (http://www.coxdocs.org/doku.php?id=maxquant:table:proteingrouptable) for column descriptions.

**Table S2. Data-frame of 6**,**427 unique proteins**. Data-frame of 6,427 rows and 194 columns, containing HUGO classification Gene Names, Entrez Protein IDs, and all Label-Free Quantitation (LFQ) intensity values.

**Table S3. Data-frame of scaled intensities for the 6**,**427 unique proteins**. Data-frame of 6,427 rows and 194 columns, containing HUGO classification Gene Names, Entrez Protein IDs, and scaled intensity values. LFQ intensities from Table S2 were grouped by protein and replicate, and scaled between 0 and 1, i.e., ‘1’ represents the fraction with the maximum intensity observed for that protein in that experiment (EXP1–EXP4), regardless of whether that maximum intensity fraction was in the Ctrl or HSP90i condition.

**Table S4. Filtered data-frame of 4**,**645 proteins found in at least 3 of 4 experiments for either treatment condition**. Data-frame of 4,645 rows and 194 columns, containing HUGO classification Gene Names, Entrez Protein IDs, and all LFQ intensity values.

**Table S5. Differential expression analysis for summed intensities from 4**,**645 filtered proteins**. Data-frame of 4,645 rows and 19 columns, containing: HUGO classification Gene Names (‘Name’); Entrez Protein IDs (‘ID’); log_2_-transformed LFQ intensities for each of the eight experiments; molecular mass in kDa (‘mw_kDa’); whether or not any intensities were imputed for the analysis (‘Imputed’); the number of missing values that needed to be imputed (‘num_NAs’); confidence intervals (CI.L and CI.R); log_2_-transformed fold change (‘LFC’); Benjamini-Hochberg-corrected adjusted *p*-values (‘p.adj’); unadjusted Student’s *t*-test *p*-values (‘p.val’); and whether or not the protein is identified as significant, based on p.adj < 0.05 and diff > 1 or < -1 (‘Significant’).

**Table S6. Characterisation of the 76 summed Differential Proteins (DPs) with respect to previous HSP90 inhibitor proteomics studies**. Data-frame of 76 rows by 17 columns, containing: HUGO classification Gene Names (‘Name’); Entrez Protein IDs (‘ID’); alternative names for protein (‘Alt_names’); unadjusted Student’s *t*-test *p*-values from DEP analysis (‘p.val’); Benjamini-Hochberg-corrected adjusted *p*-values from DEP analysis (‘p.adj’); log_2_-transformed fold change from DEP analysis (‘LFC’); whether or not the protein is identified as significant, based on p.adj < 0.05 and diff > 1 or < -1 (‘Significant’); whether the protein increased or decreased in abundance upon HSP90 inhibition (‘Direction’); whether the protein was identified as a DP in previous HSP90 proteomics studies (‘Previously_Identified_DP’), whether the protein was identified as a DP (TRUE), not identified as a DP (FALSE), or not identified at all (NA) in the specific study (‘Savitski2018’, ‘Quadroni2015’, ‘Fierro-Monti2013’, ‘Voruganti2013’, ‘Sharma2012’, ‘Wu2012’); whether or not the protein is identified as an HSP90 interactor (https://www.picard.ch/downloads/Hsp90interactors.pdf)(‘PicardList’) or an HSF1 target gene (https://hsf1base.org/)(‘HSF1_target’).

**Table S7. Differential proteins identified by PCprophet**. Data frame of 5,199 by 4 columns, containing: HUGO classification Gene Names (‘Name’); PCprophet-calculated ‘Abundance_log_marginal_likelihood’ for null and alternative hypotheses; and ‘Probability_differential_abundance’, with values > (but not equal to) 0.5 representing differential proteins.

**Table S8. HSP90i-modulated protein complexes identified by PCprophet**. Data frame of 320 by 24, containing: CO-RUM ‘ComplexID’; protein subunits from ComplexID identified as co-eluting (‘Members’); PCprophet-calculated ‘Abundance_log_marginal_likelihood’ for null and alternative hypotheses; ‘Probability_differential_abundance’, with values > (but not equal to) 0.5 representing differential complexes; CORUM ‘ComplexName’; ‘Organism’ complex was identified in; alternative names for complex (‘Synonyms’); ‘Cell line’ complex was identified in; subunits(UniProt IDs); subunits(Entrez IDs); protein complex purification method; GO description; FunCatID; FunCat description; subunits(Gene name synonyms); Complex comment; Disease comment; SWISSPROT organism; Subunits comment; gene names of all subunits annotated in the complex (‘subunits(Gene name)’); PubMed ID; full protein name of all subunits in the complex (‘subunits(Protein name)’).

**Table S9. Log**_**2**_**-transformed Fold Change (LFC) values calculated by *R* package DEP for 366 All Fraction Differential Proteins (DPs)**. Data-frame of 366 rows by 25 columns, containing: HUGO classification Gene Names (‘Name’) and log_2_-transformed fold change from DEP analysis in each fraction (‘LFC_F01’–’LFC_F24’).

**Table S10. Log**_**2**_**-transformed Fold Change (LFC) values calculated by *R* package DEP for 62 Stringent Fraction Differential Proteins (DPs)**. Data-frame of 62 rows by 28 columns, containing: HUGO classification Gene Names (‘Name’); Markov cluster ID (‘Cluster_Number’) and colour (‘Cluster_Colour’) from STRING network, as illustrated in Fig 4C; number of nodes present in the Markov cluster (‘Node_Count’); and log2-transformed fold change from DEP analysis in each fraction (‘LFC_F01’–’LFC_F24’).

## References

Armstrong, H. K., Koay, Y. C., Irani, S., Das, R., Nassar, Z. D., Australian Prostate Cancer, B., Selth, L. A., Centenera, M. M., McAlpine, S. R., and Butler, L. M. A Novel Class of Hsp90 C-Terminal Modulators Have Pre-Clinical Efficacy in Prostate Tumor Cells Without Induction of a Heat Shock Response. Prostate, 76(16):1546–1559, 2016. doi:10.1002/pros.23239.

Baas, A. F., Smit, L., and Clevers, H. LKB1 tumor suppressor protein: PARtaker in cell polarity. Trends Cell Biol, 14(6):312–9, 2004. doi:10.1016/j.tcb.2004.04.001.

Bagatell, R., Paine-Murrieta, G. D., Taylor, C. W., Pulcini, E. J., Akinaga, S., Benjamin, I. J., and Whitesell, L. Induction of a heat shock factor 1-dependent stress response alters the cytotoxic activity of hsp90-binding agents. Clin Cancer Res, 6(8):3312–8, 2000.

Bard, J. A. M., Goodall, E. A., Greene, E. R., Jonsson, E., Dong, K. C., and Martin, A. Structure and Function of the 26S Proteasome. Annu Rev Biochem, 87:697–724, 2018. doi:10.1146/annurev-biochem-062917-011931.

Benitez, M. J., Sanchez-Ponce, D., Garrido, J. J., and Wandosell, F. Hsp90 activity is necessary to acquire a proper neuronal polarization. Biochim Biophys Acta, 1843(2):245–52, 2014. doi:10.1016/j.bbamcr.2013.11.013.

Berezuk, M. A. and Schroer, T. A. Fractionation and characterization of kinesin II species in vertebrate brain. Traffic, 5(7):503–13, 2004. doi:10.1111/j.1398-9219.2004.00197.x.

Bhatia, S., Diedrich, D., Frieg, B., Ahlert, H., Stein, S., Bopp, B., Lang, F., Zang, T., Kroger, T., Ernst, T., Kogler, G., Krieg, A., Ludeke, S., Kunkel, H., Rodrigues Moita, A. J., Kassack, M. U., Marquardt, V., Opitz, F. V., Oldenburg, M., Remke, M., Babor, F., Grez, M., Hochhaus, A., Borkhardt, A., Groth, G., Nagel-Steger, L., Jose, J., Kurz, T., Gohlke, H., Hansen, F. K., and Hauer, J. Targeting HSP90 dimerization via the C terminus is effective in imatinib-resistant CML and lacks the heat shock response. Blood, 132(3):307–320, 2018. doi:10.1182/blood-2017-10-810986.

Bickel, D. and Gohlke, H. C-terminal modulators of heat shock protein of 90kDa (HSP90): State of development and modes of action. Bioorg Med Chem, 27(21):115080, 2019. doi:10.1016/j.bmc.2019.115080.

Bludau, I., Nicod, C., Martelli, C., Xue, P., Heusel, M., Fossati, A., Uliana, F., Frommelt, F., Aebersold, R., and Collins, B. C. Rapid profiling of protein complex re-organization in perturbed systems. Technical report, bioRxiv, 2021. URL https://www.biorxiv.org/content/10.1101/2021.12.17.473177v1. Section: New Results Type: article.

Brehme, M., Sverchkova, A., and Voisine, C. Proteostasis network deregulation signatures as biomarkers for pharmacological disease intervention. Curr Opin Sys Biol, 15:74–81, 2019. doi:10.1016/j.coisb.2019.03.008.

Butler, L. M., Ferraldeschi, R., Armstrong, H. K., Centenera, M. M., and Workman, P. Maximizing the Therapeutic Potential of HSP90 Inhibitors. Molecular Cancer Research, 13(11):1445–1451, 2015. doi:10.1158/1541-7786.MCR-15-0234.

Cao, P., Zhang, J., Huang, Y., Fang, Y., Lyu, J., and Shen, Y. The age-related changes and differences in energy metabolism and glutamate-glutamine recycling in the d-gal-induced and naturally occurring senescent astrocytes in vitro. Exp Gerontol, 118:9–18, 2019. doi:10.1016/j.exger.2018.12.018.

Chiosis, G. and Neckers, L. Tumor selectivity of Hsp90 inhibitors: the explanation remains elusive. ACS Chem Biol, 1(5):279–84, 2006. doi:10.1021/cb600224w.

Citri, A., Harari, D., Shohat, G., Ramakrishnan, P., Gan, J., Lavi, S., Eisenstein, M., Kimchi, A., Wallach, D., Pietrokovski, S., and Yarden, Y. Hsp90 Recognizes a Common Surface on Client Kinases. Journal of Biological Chemistry, 281(20):14361–14369, 2006. doi:10.1074/jbc.M512613200.

Conway, J. R., Lex, A., and Gehlenborg, N. UpSetR: an R package for the visualization of intersecting sets and their properties. Bioinformatics, 33(18):2938–2940, 2017. doi:10.1093/bioinformatics/btx364.

Cox, J. and Mann, M. MaxQuant enables high peptide identification rates, individualized p.p.b.-range mass accuracies and proteome-wide protein quantification. Nat Biotechnol, 26(12):1367–72, 2008. doi:10.1038/nbt.1511.

Davies, A. E. and Kaplan, K. B. Hsp90-Sgt1 and Skp1 target human Mis12 complexes to ensure efficient formation of kinetochore-microtubule binding sites. J Cell Biol, 189(2):261–74, 2010. doi:10.1083/jcb.200910036.

Donnelly, A. and Blagg, B. S. Novobiocin and additional inhibitors of the Hsp90 C-terminal nucleotide-binding pocket. Curr Med Chem, 15(26):2702–17, 2008. doi:10.2174/092986708786242895.

Dowell, J. A., Wright, L. J., Armstrong, E. A., and Denu, J. M. Benchmarking Quantitative Performance in Label-Free Proteomics. ACS Omega, 6(4):2494–2504, 2021. doi:10.1021/acsomega.0c04030.

Enright, A. J., Van Dongen, S., and Ouzounis, C. A. An efficient algorithm for large-scale detection of protein families. Nucleic Acids Res, 30(7):1575–84, 2002. doi:10.1093/nar/30.7.1575.

Eskew, J. D., Sadikot, T., Morales, P., Duren, A., Dunwiddie, I., Swink, M., Zhang, X., Hembruff, S., Donnelly, A., Rajewski, R. A., Blagg, B. S., Manjarrez, J. R., Matts, R. L., Holzbeierlein, J. M., and Vielhauer, G. A. Development and characterization of a novel C-terminal inhibitor of Hsp90 in androgen dependent and independent prostate cancer cells. BMC Cancer, 11:468, 2011. doi:10.1186/1471-2407-11-468.

Fierro-Monti, I., Echeverria, P., Racle, J., Hernandez, C., Picard, D., and Quadroni, M. Dynamic impacts of the inhibition of the molecular chaperone Hsp90 on the T-cell proteome have implications for anticancer therapy. PLoS One, 8(11):e80425, 2013. doi:10.1371/journal.pone.0080425.

Fossati, A., Li, C., Uliana, F., Wendt, F., Frommelt, F., Sykacek, P., Heusel, M., Hallal, M., Bludau, I., Capraz, T., Xue, P., Song, J., Wollscheid, B., Purcell, A. W., Gstaiger, M., and Aebersold, R. PCprophet: a framework for protein complex prediction and differential analysis using proteomic data. Nat Methods, 18(5):520–527, 2021. doi:10.1038/s41592-021-01107-5.

Fuhrmann-Stroissnigg, H., Niedernhofer, L. J., and Robbins, P. D. Hsp90 inhibitors as senolytic drugs to extend healthy aging. Cell Cycle, 17(9):1048–1055, 2018. doi:10.1080/15384101.2018.1475828.

Geladaki, A., Kocevar Britovsek, N., Breckels, L. M., Smith, T. S., Vennard, O. L., Mulvey, C. M., Crook, O. M., Gatto, L., and Lilley, K. S. Combining LOPIT with differential ultracentrifugation for high-resolution spatial proteomics. Nat Commun, 10(1):331, 2019. doi:10.1038/s41467-018-08191-w.

Gestaut, D., Roh, S. H., Ma, B., Pintilie, G., Joachimiak, L. A., Leitner, A., Walzthoeni, T., Aebersold, R., Chiu, W., and Frydman, J. The Chaperonin TRiC/CCT Associates with Prefoldin through a Conserved Electrostatic Interface Essential for Cellular Proteostasis. Cell, 177(3):751–765 e15, 2019. doi:10.1016/j.cell.2019.03.012.

Ghalhar, M. G., Akbarzadeh, A., Rahmati, M., Mellatyar, H., Dariushnejad, H., Zarghami, N., and Barkhordari, A. Comparison of inhibitory effects of 17-AAG nanoparticles and free 17-AAG on HSP90 gene expression in breast cancer. Asian Pac J Cancer Prev, 15(17):7113–8, 2014. doi:10.7314/apjcp.2014.15.17.7113.

Giulino-Roth, L., van Besien, H. J., Dalton, T., Totonchy, J. E., Rodina, A., Taldone, T., Bolaender, A., Erdjument-Bromage, H., Sadek, J., Chadburn, A., Barth, M. J., Dela Cruz, F. S., Rainey, A., Kung, A. L., Chiosis, G., and Cesarman, E. Inhibition of Hsp90 Suppresses PI3K/AKT/mTOR Signaling and Has Antitumor Activity in Burkitt Lymphoma. Mol Cancer Ther, 16(9):1779–1790, 2017. doi:10.1158/1535-7163.MCT-16-0848.

Goode, K. M., Petrov, D. P., Vickman, R. E., Crist, S. A., Pascuzzi, P. E., Ratliff, T. L., Davisson, V. J., and Hazbun, T. R. Targeting the Hsp90 C-terminal domain to induce allosteric inhibition and selective client downregulation. Biochim Biophys Acta Gen Subj, 1861(8):1992–2006, 2017. doi:10.1016/j.bbagen.2017.05.006.

Goswami, R., Russell, V. S., Tu, J. J., Thomas, C., Hughes, P., Kelly, F., Langel, S. N., Steppe, J., Palmer, S. M., Haystead, T., Blasi, M., and Permar, S. R. Oral Hsp90 inhibitor SNX-5422 attenuates SARS-CoV-2 replication and dampens inflammation in airway cells. iScience, 24(12):103412, 2021. doi:10.1016/j.isci.2021.103412.

Hall, P. A., Todd, C. B., Hyland, P. L., McDade, S. S., Grabsch, H., Dattani, M., Hillan, K. J., and Russell, S. E. The septin-binding protein anillin is overexpressed in diverse human tumors. Clin Cancer Res, 11(19 Pt 1):6780–6, 2005. doi:10.1158/1078-0432.CCR-05-0997.

Hanahan, D. and Weinberg, R. A. Hallmarks of cancer: the next generation. Cell, 144(5):646–74, 2011. doi:10.1016/j.cell.2011.02.013.

Hartson, S. D. and Matts, R. L. Approaches for defining the Hsp90-dependent proteome. Biochim Biophys Acta, 1823(3):656–67, 2012. doi:10.1016/j.bbamcr.2011.08.013.

Hasan, A., Haque, E., Hameed, R., Maier, P. N., Irfan, S., Kamil, M., Nazir, A., and Mir, S. S. Hsp90 inhibitor gedunin causes apoptosis in A549 lung cancer cells by disrupting Hsp90:Beclin-1:Bcl-2 interaction and downregulating autophagy. Life Sci, 256:118000, 2020. doi:10.1016/j.lfs.2020.118000.

Haslbeck, M., Weinkauf, S., and Buchner, J. Small heat shock proteins: Simplicity meets complexity. J Biol Chem, 294(6):2121–2132, 2019. doi:10.1074/jbc.REV118.002809.

Heusel, M., Bludau, I., Rosenberger, G., Hafen, R., Frank, M., Banaei-Esfahani, A., van Drogen, A., Collins, B. C., Gstaiger, M., and Aebersold, R. Complex-centric proteome profiling by SEC-SWATH-MS. Mol Syst Biol, 15(1):e8438, 2019. doi:10.15252/msb.20188438.

Heusel, M., Frank, M., Kohler, M., Amon, S., Frommelt, F., Rosenberger, G., Bludau, I., Aulakh, S., Linder, M. I., Liu, Y., Collins, B. C., Gstaiger, M., Kutay, U., and Aebersold, R. A Global Screen for Assembly State Changes of the Mitotic Proteome by SEC-SWATH-MS. Cell Syst, 2020. doi:10.1016/j.cels.2020.01.001.

Hipp, M. S., Kasturi, P., and Hartl, F. U. The proteostasis network and its decline in ageing. Nat Rev Mol Cell Biol, 20(7):421–435, 2019. doi:10.1038/s41580-019-0101-y.

Holford, J., Sharp, S. Y., Murrer, B. A., Abrams, M., and Kelland, L. R. In vitro circumvention of cisplatin resistance by the novel sterically hindered platinum complex AMD473. British Journal of Cancer, 77 (3):366–373, 1998. doi:10.1038/bjc.1998.59.

Honma, Y., Kurokawa, Y., Sawaki, A., Naito, Y., Iwagami, S., Baba, H., Komatsu, Y., Nishida, T., and Doi, T. Randomized, double-blind, placebo (PL)-controlled, phase III trial of pimitespib (TAS-116), an oral inhibitor of heat shock protein 90 (HSP90), in patients (pts) with advanced gastrointestinal stromal tumor (GIST) refractory to imatinib (IM), sunitinib (SU) and regorafenib (REG). Journal of Clinical Oncology, 39(15_suppl):11524–11524, 2021. doi:10.1200/JCO.2021.39.15_suppl.11524.

Huh, T. L., Kim, Y. O., Ko, H. J., Kim, S. H., Park, H. C., Lee, L. J., and Song, B. J. Human mitochondrial NAD-specific isocitrate dehrydrogenase 3 and subunits: Cloning, tissue specific expression and functional analyses of the recombinant proteins. FASEB Journal, 11(9), 1997.

Itzhak, D. N., Tyanova, S., Cox, J., and Borner, G. H. Global, quantitative and dynamic mapping of protein subcellular localization. Elife, 5, 2016. doi:10.7554/eLife.16950.

Jaeger, A. M. and Whitesell, L. HSP90: Enabler of Cancer Adaptation. Annual Review of Cancer Biology, 3(1):275–297, 2019. doi:10.1146/annurev-cancerbio-030518-055533.

Janssens, G. E., Lin, X.-X., Millan-Ariño, L., Kavšek, A., Sen, I., Seinstra, R. I., Stroustrup, N., Nollen, E. A. A., and Riedel, C. G. Transcriptomics-Based Screening Identifies Pharmacological Inhibition of Hsp90 as a Means to Defer Aging. Cell Reports, 27(2):467–480.e6, 2019. doi:10.1016/j.celrep.2019.03.044.

Jin, L., Bi, Y., Hu, C., Qu, J., Shen, S., Wang, X., and Tian, Y. A comparative study of evaluating missing value imputation methods in label-free proteomics. Sci Rep, 11(1):1760, 2021. doi:10.1038/s41598-021-81279-4.

Kamal, A., Thao, L., Sensintaffar, J., Zhang, L., Boehm, M. F., Fritz, L. C., and Burrows, F. J. A high-affinity conformation of Hsp90 confers tumour selectivity on Hsp90 inhibitors. Nature, 425(6956): 407–10, 2003. doi:10.1038/nature01913.

Karkoulis, P. K., Stravopodis, D. J., Margaritis, L. H., and Voutsinas, G. E. 17-Allylamino-17-demethoxygeldanamycin induces downregulation of critical Hsp90 protein clients and results in cell cycle arrest and apoptosis of human urinary bladder cancer cells. BMC Cancer, 10:481, 2010. doi:10.1186/1471-2407-10-481.

Khandelwal, A., Kent, C. N., Balch, M., Peng, S., Mishra, S. J., Deng, J., Day, V. W., Liu, W., Subramanian, C., Cohen, M., Holzbeierlein, J. M., Matts, R., and Blagg, B. S. J. Structure-guided design of an Hsp90beta N-terminal isoform-selective inhibitor. Nat Commun, 9(1):425, 2018. doi:10.1038/s41467-017-02013-1.

Kirkwood, K. J., Ahmad, Y., Larance, M., and Lamond, A. I. Characterization of native protein complexes and protein isoform variation using size-fractionation-based quantitative proteomics. Mol Cell Proteomics, 12(12):3851–73, 2013. doi:10.1074/mcp.M113.032367.

Kovacs, D., Sigmond, T., Hotzi, B., Bohar, B., Fazekas, D., Deak, V., Vellai, T., and Barna, J. HSF1Base: A Comprehensive Database of HSF1 (Heat Shock Factor 1) Target Genes. Int J Mol Sci, 20(22), 2019. doi:10.3390/ijms20225815.

Lacey, T. and Lacey, H. Linking hsp90’s role as an evolutionary capacitator to the development of cancer. Cancer Treatment and Research Communications, 28:100400, 2021. doi:10.1016/j.ctarc.2021.100400.

Landry, J. J., Pyl, P. T., Rausch, T., Zichner, T., Tekkedil, M. M., Stutz, A. M., Jauch, A., Aiyar, R. S., Pau, G., Delhomme, N., Gagneur, J., Korbel, J. O., Huber, W., and Steinmetz, L. M. The genomic and transcriptomic landscape of a HeLa cell line. G3 (Bethesda), 3(8):1213–24, 2013. doi:10.1534/g3.113.005777.

Lange, B. M., Rebollo, E., Herold, A., and Gonzalez, C. Cdc37 is essential for chromosome segregation and cytokinesis in higher eukaryotes. EMBO J, 21(20):5364–74, 2002. doi:10.1093/emboj/cdf531.

Larance, M., Ahmad, Y., Kirkwood, K. J., Ly, T., and Lamond, A. I. Global subcellular characterization of protein degradation using quantitative proteomics. Mol Cell Proteomics, 12(3):638–50, 2013. doi:10.1074/mcp.M112.024547.

Larance, M., Kirkwood, K. J., Tinti, M., Brenes Murillo, A., Ferguson, M. A., and Lamond, A. I. Global Membrane Protein Interactome Analysis using In vivo Crosslinking and Mass Spectrometry-based Protein Correlation Profiling. Mol Cell Proteomics, 15(7):2476–90, 2016. doi:10.1074/mcp.O115.055467.

Li, L., Chen, N., Xia, D., Xu, S., Dai, W., Tong, Y., Wang, L., Jiang, Z., You, Q., and Xu, X. Discovery of a covalent inhibitor of heat shock protein 90 with antitumor activity that blocks the co-chaperone binding via C-terminal modification. Cell Chem Biol, 28(10):1446–1459 e6, 2021. doi:10.1016/j.chembiol.2021.03.016.

Lian, Y. F., Huang, Y. L., Wang, J. L., Deng, M. H., Xia, T. L., Zeng, M. S., Chen, M. S., Wang, H. B., and Huang, Y. H. Anillin is required for tumor growth and regulated by miR-15a/miR-16-1 in HBV-related hepatocellular carcinoma. Aging (Albany NY), 10(8):1884–1901, 2018. doi:10.18632/aging.101510.

Lin, Y. C., Boone, M., Meuris, L., Lemmens, I., Van Roy, N., Soete, A., Reumers, J., Moisse, M., Plaisance, S., Drmanac, R., Chen, J., Speleman, F., Lambrechts, D., Van de Peer, Y., Tavernier, J., and Callewaert, N. Genome dynamics of the human embryonic kidney 293 lineage in response to cell biology manipulations. Nat Commun, 5:4767, 2014. doi:10.1038/ncomms5767.

Liu, X. Y., Seh, C. C., and Cheung, P. C. HSP90 is required for TAK1 stability but not for its activation in the pro-inflammatory signaling pathway. FEBS Lett, 582(29):4023–31, 2008. doi:10.1016/j.febslet.2008.10.053.

Ludwig, C., Gillet, L., Rosenberger, G., Amon, S., Collins, B. C., and Aebersold, R. Data-independent acquisition-based SWATH-MS for quantitative proteomics: a tutorial. Mol Syst Biol, 14(8):e8126, 2018. doi:10.15252/msb.20178126.

Makhnevych, T. and Houry, W. A. The role of Hsp90 in protein complex assembly. Biochimica Et Biophysica Acta, 1823(3):674–682, 2012. doi:10.1016/j.bbamcr.2011.09.001.

Maloney, A., Clarke, P. A., Naaby-Hansen, S., Stein, R., Koopman, J. O., Akpan, A., Yang, A., Zvelebil, M., Cramer, R., Stimson, L., Aherne, W., Banerji, U., Judson, I., Sharp, S., Powers, M., deBilly, E., Salmons, J., Walton, M., Burlingame, A., Waterfield, M., and Workman, P. Gene and protein expression profiling of human ovarian cancer cells treated with the heat shock protein 90 inhibitor 17-allylamino-17-demethoxygeldanamycin. Cancer Res, 67(7):3239–53, 2007. doi:10.1158/0008-5472.CAN-06-2968.

Masser, A. E., Ciccarelli, M., and Andréasson, C. Hsf1 on a leash - controlling the heat shock response by chaperone titration. Experimental Cell Research, 396(1):112246, 2020. doi:10.1016/j.yexcr.2020.112246.

May, J. L., Kouri, F. M., Hurley, L. A., Liu, J., Tommasini-Ghelfi, S., Ji, Y., Gao, P., Calvert, A. E., Lee, A., Chandel, N. S., Davuluri, R. V., Horbinski, C. M., Locasale, J. W., and Stegh, A. H. IDH3alpha regulates one-carbon metabolism in glioblastoma. Sci Adv, 5(1):eaat0456, 2019. doi:10.1126/sciadv.aat0456.

McKinley, K. L. and Cheeseman, I. M. Polo-like kinase 1 licenses CENP-A deposition at centromeres. Cell, 158(2):397–411, 2014. doi:10.1016/j.cell.2014.06.016.

Miao, W., Li, L., Zhao, Y., Dai, X., Chen, X., and Wang, Y. HSP90 inhibitors stimulate DNAJB4 protein expression through a mechanism involving N(6)-methyladenosine. Nat Commun, 10(1):3613, 2019. doi:10.1038/s41467-019-11552-8.

Mimnaugh, E. G., Chavany, C., and Neckers, L. Polyubiquitination and proteasomal degradation of the p185c-erbB-2 receptor protein-tyrosine kinase induced by geldanamycin. J Biol Chem, 271(37): 22796–801, 1996. doi:10.1074/jbc.271.37.22796.

Mitsopoulos, C., Di Micco, P., Fernandez, E. V., Dolciami, D., Holt, E., Mica, I. L., Coker, E. A., Tym, J. E., Campbell, J., Che, K. H., Ozer, B., Kannas, C., Antolin, A. A., Workman, P., and Al-Lazikani, B. canSAR: update to the cancer translational research and drug discovery knowledgebase. Nucleic Acids Research, 49(D1):D1074–D1082, 2021. doi:10.1093/nar/gkaa1059.

Moulick, K., Ahn, J. H., Zong, H., Rodina, A., Cerchietti, L., Gomes DaGama, E. M., Caldas-Lopes, E., Beebe, K., Perna, F., Hatzi, K., Vu, L. P., Zhao, X., Zatorska, D., Taldone, T., Smith-Jones, P., Alpaugh, M., Gross, S. S., Pillarsetty, N., Ku, T., Lewis, J. S., Larson, S. M., Levine, R., Erdjument-Bromage, H., Guzman, M. L., Nimer, S. D., Melnick, A., Neckers, L., and Chiosis, G. Affinity-based proteomics reveal cancer-specific networks coordinated by Hsp90. Nat Chem Biol, 7(11):818–26, 2011. doi:10.1038/nchembio.670.

Moullintraffort, L., Bruneaux, M., Nazabal, A., Allegro, D., Giudice, E., Zal, F., Peyrot, V., Barbier, P., Thomas, D., and Garnier, C. Biochemical and biophysical characterization of the Mg2+-induced 90-kDa heat shock protein oligomers. J Biol Chem, 285(20):15100–15110, 2010. doi:10.1074/jbc.M109.094698.

Mulvey, C. M., Breckels, L. M., Geladaki, A., Britovsek, N. K., Nightingale, D. J. H., Christoforou, A., Elzek, M., Deery, M. J., Gatto, L., and Lilley, K. S. Using hyperLOPIT to perform high-resolution mapping of the spatial proteome. Nat Protoc, 12(6):1110–1135, 2017. doi:10.1038/nprot.2017.026.

Muntel, J., Kirkpatrick, J., Bruderer, R., Huang, T., Vitek, O., Ori, A., and Reiter, L. Comparison of Protein Quantification in a Complex Background by DIA and TMT Workflows with Fixed Instrument Time. J Proteome Res, 18(3):1340–1351, 2019. doi:10.1021/acs.jproteome.8b00898.

Naydenov, N. G., Koblinski, J. E., and Ivanov, A. I. Anillin is an emerging regulator of tumorigenesis, acting as a cortical cytoskeletal scaffold and a nuclear modulator of cancer cell differentiation. Cell Mol Life Sci, 78(2):621–633, 2021. doi:10.1007/s00018-020-03605-9.

Niikura, Y., Ohta, S., Vandenbeldt, K. J., Abdulle, R., McEwen, B. F., and Kitagawa, K. 17-AAG, an Hsp90 inhibitor, causes kinetochore defects: a novel mechanism by which 17-AAG inhibits cell proliferation. Oncogene, 25(30):4133–46, 2006. doi:10.1038/sj.onc.1209461.

Niikura, Y., Kitagawa, R., Ogi, H., and Kitagawa, K. SGT1-HSP90 complex is required for CENP-A deposition at centromeres. Cell Cycle, 16(18):1683–1694, 2017. doi:10.1080/15384101.2017.1325039.

O’Connell, J. D., Paulo, J. A., O’Brien, J. J., and Gygi, S. P. Proteome-Wide Evaluation of Two Common Protein Quantification Methods. J Proteome Res, 17(5):1934–1942, 2018. doi:10.1021/acs.jproteome.8b00016.

Olivoto, T. and Lúcio, A. D. metan: An R package for multi-environment trial analysis. Methods in Ecology and Evolution, 11(6):783–789, 2020. doi:10.1111/2041-210X.13384.

Pang, C. N. I., Ballouz, S., Weissberger, D., Thibaut, L. M., Hamey, J. J., Gillis, J., Wilkins, M. R., and Hart-Smith, G. Analytical Guidelines for co-fractionation Mass Spectrometry Obtained through Global Profiling of Gold Standard Saccharomyces cerevisiae Protein Complexes. Molecular & cellular proteomics: MCP, 19(11):1876–1895, 2020. doi:10.1074/mcp.RA120.002154.

Park, J. M., Kim, Y. J., Park, S., Park, M., Farrand, L., Nguyen, C. T., Ann, J., Nam, G., Park, H. J., Lee, J., Kim, J. Y., and Seo, J. H. A novel HSP90 inhibitor targeting the C-terminal domain attenuates trastuzumab resistance in HER2-positive breast cancer. Mol Cancer, 19(1):161, 2020. doi:10.1186/s12943-020-01283-6.

Park, H. K., Yoon, N. G., Lee, J. E., Hu, S., Yoon, S., Kim, S. Y., Hong, J. H., Nam, D., Chae, Y. C., Park, J. B., and Kang, B. H. Unleashing the full potential of Hsp90 inhibitors as cancer therapeutics through simultaneous inactivation of Hsp90, Grp94, and TRAP1. Exp Mol Med, 52(1):79–91, 2020. doi:10.1038/s12276-019-0360-x.

Pearl, L. H. Review: The HSP90 molecular chaperone-an enigmatic ATPase. Biopolymers, 105(8): 594–607, 2016. doi:10.1002/bip.22835.

Perez-Riverol, Y., Bai, J., Bandla, C., García-Seisdedos, D., Hewapathirana, S., Kamatchinathan, S., Kundu, D., Prakash, A., Frericks-Zipper, A., Eisenacher, M., Walzer, M., Wang, S., Brazma, A., and Vizcaíno, J. The PRIDE database resources in 2022: a hub for mass spectrometry-based proteomics evidences. Nucleic Acids Research, (gkab1038), 2021. doi:10.1093/nar/gkab1038.

Piekny, A. J. and Maddox, A. S. The myriad roles of Anillin during cytokinesis. Semin Cell Dev Biol, 21 (9):881–91, 2010. doi:10.1016/j.semcdb.2010.08.002.

Pincus, D. Regulation of Hsf1 and the Heat Shock Response. Advances in Experimental Medicine and Biology, 1243:41–50, 2020. doi:10.1007/978-3-030-40204-4_3.

Pomaznoy, M., Ha, B., and Peters, B. GOnet: a tool for interactive Gene Ontology analysis. BMC Bioinformatics, 19(1):470, 2018. doi:10.1186/s12859-018-2533-3.

Quadroni, M., Potts, A., and Waridel, P. Hsp90 inhibition induces both protein-specific and global changes in the ubiquitinome. J Proteomics, 120:215–29, 2015. doi:10.1016/j.jprot.2015.02.020.

Rodina, A., Wang, T., Yan, P., Gomes, E. D., Dunphy, M. P., Pillarsetty, N., Koren, J., Gerecitano, J. F., Taldone, T., Zong, H., Caldas-Lopes, E., Alpaugh, M., Corben, A., Riolo, M., Beattie, B., Pressl, C., Peter, R. I., Xu, C., Trondl, R., Patel, H. J., Shimizu, F., Bolaender, A., Yang, C., Panchal, P., Farooq, M. F., Kishinevsky, S., Modi, S., Lin, O., Chu, F., Patil, S., Erdjument-Bromage, H., Zanzonico, P., Hudis, C., Studer, L., Roboz, G. J., Cesarman, E., Cerchietti, L., Levine, R., Melnick, A., Larson, S. M., Lewis, J. S., Guzman, M. L., and Chiosis, G. The epichaperome is an integrated chaperome network that facilitates tumour survival. Nature, 538(7625):397–401, 2016. doi:10.1038/nature19807.

Ruepp, A., Waegele, B., Lechner, M., Brauner, B., Dunger-Kaltenbach, I., Fobo, G., Frishman, G., Montrone, C., and Mewes, H. W. CORUM: the comprehensive resource of mammalian protein complexes–2009. Nucleic Acids Res, 38(Database issue):D497–501, 2010. doi:10.1093/nar/gkp914.

Salas, D., Stacey, R. G., Akinlaja, M., and Foster, L. J. Next-generation Interactomics: Considerations for the Use of Co-elution to Measure Protein Interaction Networks. Mol Cell Proteomics, 19(1):1–10, 2020. doi:10.1074/mcp.R119.001803.

Samant, R. S., Clarke, P. A., and Workman, P. The expanding proteome of the molecular chaperone HSP90. Cell Cycle, 11(7):1301–8, 2012. doi:10.4161/cc.19722.

Samant, R. S., Clarke, P. A., and Workman, P. E3 ubiquitin ligase Cullin-5 modulates multiple molecular and cellular responses to heat shock protein 90 inhibition in human cancer cells. Proc Natl Acad Sci U S A, 111(18):6834–9, 2014. doi:10.1073/pnas.1322412111.

Sanchez-Martin, C., Moroni, E., Ferraro, M., Laquatra, C., Cannino, G., Masgras, I., Negro, A., Quadrelli, P., Rasola, A., and Colombo, G. Rational Design of Allosteric and Selective Inhibitors of the Molecular Chaperone TRAP1. Cell Rep, 31(3):107531, 2020. doi:10.1016/j.celrep.2020.107531.

Savitski, M. M., Zinn, N., Faelth-Savitski, M., Poeckel, D., Gade, S., Becher, I., Muelbaier, M., Wagner, J., Strohmer, K., Werner, T., Melchert, S., Petretich, M., Rutkowska, A., Vappiani, J., Franken, H., Steidel, M., Sweetman, G. M., Gilan, O., Lam, E. Y. N., Dawson, M. A., Prinjha, R. K., Grandi, P., Bergamini, G., and Bantscheff, M. Multiplexed Proteome Dynamics Profiling Reveals Mechanisms Controlling Protein Homeostasis. Cell, 173(1):260–274 e25, 2018. doi:10.1016/j.cell.2018.02.030.

Schlossarek, D., Luzarowski, M., Sokolowska, E., Górka, M., Willmitzer, L., and Skirycz, A. PROMISed: A novel web-based tool to facilitate analysis and visualization of the molecular interaction networks from co-fractionation mass spectrometry (CF-MS) experiments. Computational and structural biotechnology journal, 19:5117–5125, 2021. doi:10.1016/j.csbj.2021.08.042.

Schopf, F. H., Biebl, M. M., and Buchner, J. The HSP90 chaperone machinery. Nat Rev Mol Cell Biol, 18 (6):345–360, 2017. doi:10.1038/nrm.2017.20.

Schulte, T. W., Blagosklonny, M. V., Ingui, C., and Neckers, L. Disruption of the Raf-1-Hsp90 molecular complex results in destabilization of Raf-1 and loss of Raf-1-Ras association. J Biol Chem, 270(41): 24585–8, 1995. doi:10.1074/jbc.270.41.24585.

Schulte, T. W., Blagosklonny, M. V., Romanova, L., Mushinski, J. F., Monia, B. P., Johnston, J. F., Nguyen, P., Trepel, J., and Neckers, L. M. Destabilization of Raf-1 by geldanamycin leads to disruption of the Raf-1-MEK-mitogen-activated protein kinase signalling pathway. Mol Cell Biol, 16(10):5839–45, 1996. doi:10.1128/mcb.16.10.5839.

Schumacher, J. A., Crockett, D. K., Elenitoba-Johnson, K. S. J., and Lim, M. S. Proteome-wide changes induced by the Hsp90 inhibitor, geldanamycin in anaplastic large cell lymphoma cells. PROTEOMICS, 7(15):2603–2616, 2007. doi:10.1002/pmic.200700108.

Serwetnyk, M. A. and Blagg, B. S. J. The disruption of protein-protein interactions with co-chaperones and client substrates as a strategy towards Hsp90 inhibition. Acta Pharm Sin B, 11(6):1446–1468, 2021. doi:10.1016/j.apsb.2020.11.015.

Sharma, K., Vabulas, R. M., Macek, B., Pinkert, S., Cox, J., Mann, M., and Hartl, F. U. Quantitative proteomics reveals that Hsp90 inhibition preferentially targets kinases and the DNA damage response. Mol Cell Proteomics, 11(3):M111 014654, 2012. doi:10.1074/mcp.M111.014654.

Shi, L., Zhang, Z., Fang, S., Xu, J., Liu, J., Shen, J., Fang, F., Luo, L., and Yin, Z. Heat shock protein 90 (Hsp90) regulates the stability of transforming growth factor beta-activated kinase 1 (TAK1) in interleukin-1beta-induced cell signaling. Mol Immunol, 46(4):541–50, 2009. doi:10.1016/j.molimm.2008.07.019.

Soroka, J., Wandinger, S. K., Mausbacher, N., Schreiber, T., Richter, K., Daub, H., and Buchner, J. Conformational switching of the molecular chaperone Hsp90 via regulated phosphorylation. Mol Cell, 45(4):517–28, 2012. doi:10.1016/j.molcel.2011.12.031.

Stepath, M., Zulch, B., Maghnouj, A., Schork, K., Turewicz, M., Eisenacher, M., Hahn, S., Sitek, B., and Bracht, T. Systematic Comparison of Label-Free, SILAC, and TMT Techniques to Study Early Adaption toward Inhibition of EGFR Signaling in the Colorectal Cancer Cell Line DiFi. J Proteome Res, 19(2):926–937, 2020. doi:10.1021/acs.jproteome.9b00701.

Stothert, A. R., Suntharalingam, A., Tang, X., Crowley, V. M., Mishra, S. J., Webster, J. M., Nordhues, 2. A., Huard, D. J. E., Passaglia, C. L., Lieberman, R. L., Blagg, B. S. J., Blair, L. J., Koren, J., 3rd, and Dickey, C. A. Isoform-selective Hsp90 inhibition rescues model of hereditary open-angle glaucoma. Sci Rep, 7(1):17951, 2017. doi:10.1038/s41598-017-18344-4.

Sui, X., Pires, D. E. V., Ormsby, A. R., Cox, D., Nie, S., Vecchi, G., Vendruscolo, M., Ascher, D. B., Reid, G. E., and Hatters, D. M. Widespread remodeling of proteome solubility in response to different protein homeostasis stresses. Proc Natl Acad Sci U S A, 117(5):2422–2431, 2020. doi:10.1073/pnas.1912897117.

Szklarczyk, D., Gable, A. L., Nastou, K. C., Lyon, D., Kirsch, R., Pyysalo, S., Doncheva, N. T., Legeay, M., Fang, T., Bork, P., Jensen, L. J., and von Mering, C. The STRING database in 2021: customizable protein-protein networks, and functional characterization of user-uploaded gene/measurement sets. Nucleic Acids Res, 49(D1):D605–D612, 2021. doi:10.1093/nar/gkaa1074.

Taipale, M., Krykbaeva, I., Koeva, M., Kayatekin, C., Westover, K., Karras, G., and Lindquist, S. Quantitative Analysis of Hsp90-Client Interactions Reveals Principles of Substrate Recognition. Cell, 150 (5):987–1001, 2012. doi:10.1016/j.cell.2012.06.047.

Terasawa, K. and Minami, Y. A client-binding site of Cdc37. FEBS J, 272(18):4684–90, 2005. doi:10.1111/j.1742-4658.2005.04884.x.

Terracciano, S., Russo, A., Chini, M. G., Vaccaro, M. C., Potenza, M., Vassallo, A., Riccio, R., Bifulco, G., and Bruno, I. Discovery of new molecular entities able to strongly interfere with Hsp90 C-terminal domain. Sci Rep, 8(1):1709, 2018. doi:10.1038/s41598-017-14902-y.

Tommasini-Ghelfi, S., Murnan, K., Kouri, F. M., Mahajan, A. S., May, J. L., and Stegh, A. H. Cancer-associated mutation and beyond: The emerging biology of isocitrate dehydrogenases in human disease. Sci Adv, 5(5):eaaw4543, 2019. doi:10.1126/sciadv.aaw4543.

Tuan, N. M. and Lee, C. H. Role of Anillin in Tumour: From a Prognostic Biomarker to a Novel Target. Cancers (Basel), 12(6), 2020. doi:10.3390/cancers12061600.

Tyanova, S., Temu, T., and Cox, J. The MaxQuant computational platform for mass spectrometry-based shotgun proteomics. Nat Protoc, 11(12):2301–2319, 2016. doi:10.1038/nprot.2016.136.

Vaughan, C. K., Mollapour, M., Smith, J. R., Truman, A., Hu, B., Good, V. M., Panaretou, B., Neckers, L., Clarke, P. A., Workman, P., Piper, P. W., Prodromou, C., and Pearl, L. H. Hsp90-Dependent Activation of Protein Kinases Is Regulated by Chaperone-Targeted Dephosphorylation of Cdc37. Molecular Cell, 31(6):886–895, 2008. doi:10.1016/j.molcel.2008.07.021.

Voruganti, S., Lacroix, J. C., Rogers, C. N., Rogers, J., Matts, R. L., and Hartson, S. D. The anticancer drug AUY922 generates a proteomics fingerprint that is highly conserved among structurally diverse Hsp90 inhibitors. J Proteome Res, 12(8):3697–706, 2013. doi:10.1021/pr400321x.

Wang, G., Shen, W., Cui, L., Chen, W., Hu, X., and Fu, J. Overexpression of Anillin (ANLN) is correlated with colorectal cancer progression and poor prognosis. Cancer Biomark, 16(3):459–65, 2016. doi:10.3233/CBM-160585.

Wang, Y., Jin, F., Wang, R., Li, F., Wu, Y., Kitazato, K., and Wang, Y. HSP90: a promising broad-spectrum antiviral drug target. Archives of Virology, 162(11):3269–3282, 2017. doi:10.1007/s00705-017-3511-1.

Wang, T., Rodina, A., Dunphy, M. P., Corben, A., Modi, S., Guzman, M. L., Gewirth, D. T., and Chiosis, G. Chaperome heterogeneity and its implications for cancer study and treatment. J Biol Chem, 294 (6):2162–2179, 2019. doi:10.1074/jbc.REV118.002811.

Webb-Robertson, B. J., Wiberg, H. K., Matzke, M. M., Brown, J. N., Wang, J., McDermott, J. E., Smith, R. D., Rodland, K. D., Metz, T. O., Pounds, J. G., and Waters, K. M. Review, evaluation, and discussion of the challenges of missing value imputation for mass spectrometry-based label-free global proteomics. J Proteome Res, 14(5):1993–2001, 2015. doi:10.1021/pr501138h.

Weidenauer, L., Wang, T., Joshi, S., Chiosis, G., and Quadroni, M. R. Proteomic interrogation of HSP90 and insights for medical research. Expert Rev Proteomics, 14(12):1105–1117, 2017. doi:10.1080/14789450.2017.1389649.

Workman, P. Altered states: selectively drugging the Hsp90 cancer chaperone. Trends Mol Med, 10(2): 47–51, 2004. doi:10.1016/j.molmed.2003.12.005.

Workman, P. Reflections and Outlook on Targeting HSP90, HSP70 and HSF1 in Cancer: A Personal Perspective. Advances in Experimental Medicine and Biology, 1243:163–179, 2020. doi:10.1007/978-3-030-40204-4_11.

Wu, Z., Gholami, A. M., and Kuster, B. Systematic identification of the HSP90 candidate regulated proteome. Mol Cell Proteomics, 11(6):M111 016675, 2012. doi:10.1074/mcp.M111.016675.

Xu, C., Liu, J., Hsu, L. C., Luo, Y., Xiang, R., and Chuang, T. H. Functional interaction of heat shock protein 90 and Beclin 1 modulates Toll-like receptor-mediated autophagy. FASEB J, 25(8):2700–10, 2011. doi:10.1096/fj.10-167676.

Xu, J., Zheng, H., Yuan, S., Zhou, B., Zhao, W., Pan, Y., and Qi, D. Overexpression of ANLN in lung adenocarcinoma is associated with metastasis. Thorac Cancer, 10(8):1702–1709, 2019. doi:10.1111/1759-7714.13135.

Yuno, A., Lee, M. J., Lee, S., Tomita, Y., Rekhtman, D., Moore, B., and Trepel, J. B. Clinical Evaluation and Biomarker Profiling of Hsp90 Inhibitors. Methods Mol Biol, 1709:423–441, 2018. doi:10.1007/978-1-4939-7477-1_29.

Zavareh, R. B., Spangenberg, S. H., Woods, A., Martínez-Peña, F., and Lairson, L. L. HSP90 Inhibition Enhances Cancer Immunotherapy by Modulating the Surface Expression of Multiple Immune Check-point Proteins. Cell Chemical Biology, 28(2):158–168.e5, 2021. doi:10.1016/j.chembiol.2020.10.005.

Zeng, L., Morinibu, A., Kobayashi, M., Zhu, Y., Wang, X., Goto, Y., Yeom, C. J., Zhao, T., Hirota, K., Shinomiya, K., Itasaka, S., Yoshimura, M., Guo, G., Hammond, E. M., Hiraoka, M., and Harada, H. Aberrant IDH3alpha expression promotes malignant tumor growth by inducing HIF-1-mediated metabolic reprogramming and angiogenesis. Oncogene, 34(36):4758–66, 2015. doi:10.1038/onc.2014.411.

Zhang, D., Wang, Y., Shi, Z., Liu, J., Sun, P., Hou, X., Zhang, J., Zhao, S., Zhou, B. P., and Mi, J. Metabolic reprogramming of cancer-associated fibroblasts by IDH3alpha downregulation. Cell Rep, 10(8):1335–48, 2015. doi:10.1016/j.celrep.2015.02.006.

Zhang, X., Smits, A. H., van Tilburg, G. B., Ovaa, H., Huber, W., and Vermeulen, M. Proteome-wide identification of ubiquitin interactions using UbIA-MS. Nat Protoc, 13(3):530–550, 2018. doi:10.1038/nprot.2017.147.

Zhang, S., Nguyen, L. H., Zhou, K., Tu, H. C., Sehgal, A., Nassour, I., Li, L., Gopal, P., Goodman, J., Singal, A. G., Yopp, A., Zhang, Y., Siegwart, D. J., and Zhu, H. Knockdown of Anillin Actin Binding Protein Blocks Cytokinesis in Hepatocytes and Reduces Liver Tumor Development in Mice Without Affecting Regeneration. Gastroenterology, 154(5):1421–1434, 2018a. doi:10.1053/j.gastro.2017.12.013.

Zhang, J., Liu, X., Yu, G., Liu, L., Wang, J., Chen, X., Bian, Y., Ji, Y., Zhou, X., Chen, Y., Ji, J., Xiang, Z., Guo, L., Fang, J., Sun, Y., Cao, H., Zhu, Z., and Yu, Y. UBE2C Is a Potential Biomarker of Intestinal-Type Gastric Cancer With Chromosomal Instability. Front Pharmacol, 9:847, 2018b. doi:10.3389/fphar.2018.00847.

Zhu, D., Li, S., Chen, C., Wang, S., Zhu, J., Kong, L., and Luo, J. Tubocapsenolide A targets C-terminal cysteine residues of HSP90 to exert the anti-tumor effect. Pharmacol Res, 166:105523, 2021. doi:10.1016/j.phrs.2021.105523.

